# Neurotransmitter Classification from Electron Microscopy Images at Synaptic Sites in Drosophila Melanogaster

**DOI:** 10.1101/2020.06.12.148775

**Authors:** Nils Eckstein, Alexander Shakeel Bates, Andrew Champion, Michelle Du, Yijie Yin, Philipp Schlegel, Alicia Kun-Yang Lu, Thomson Rymer, Samantha Finley-May, Tyler Paterson, Ruchi Parekh, Sven Dorkenwald, Arie Matsliah, Szi-Chieh Yu, Claire McKellar, Amy Sterling, Katharina Eichler, Marta Costa, Sebastian Seung, Mala Murthy, Volker Hartenstein, Gregory S.X.E. Jefferis, Jan Funke

## Abstract

High-resolution electron microscopy of nervous systems enables the reconstruction of connectomes. A key piece of missing information from connectomes is the synaptic sign. We show that for *D. melanogaster*, artificial neural networks can predict the transmitter type released at synapses from electron micrographs and thus add putative signs to connections. Our network discriminates between six transmitters (acetylcholine, glutamate, GABA, serotonin, dopamine, octopamine) with an average accuracy of 87%/94% for synapses/entire neurons. We developed an explainability method to reveal which features our network is using and found significant ultrastructural differences between the classical transmitters. We predict transmitters in two connectomes and characterize morphological and connection properties of tens of thousands of neurons classed by predicted transmitter expression. We find that hemilineages in *D. melanogaster* largely express only one fast-acting transmitter among their neurons. Furthermore, we show that neurons with different transmitters may differ in features like polarization and projection targets.

## 1 Introduction

Generating a connectome entails identifying all neurons and the synapses that connect them. In recent years, advances in imaging technology have enabled high-resolution electron microscopy (EM) imaging of whole brain or nervous system data sets (Zheng et al., 2018; Ryan et al., 2016; Cook et al., 2019; Ohyama et al., 2015; Phelps et al., 2021), opening up the possibility of generating neuron-level wiring diagrams (“connectomes”) of entire nervous systems. In addition, recent advances in automated methods for segmenting neurons (Funke et al., 2018; Januszewski et al., 2018; Lee et al., 2019; Sheridan et al., 2022), detecting synapses (Kreshuk et al., 2015; Staffler et al., 2017; Buhmann et al., 2019; Huang et al., 2018) and proofreading (Dorkenwald et al., 2022) greatly reduce the time of human involvement in these tasks. They have recently been applied to generate the connectome for a large part of the *D. melanogaster* brain (Scheffer et al., 2020a) and are being used to generate the first full brain connectome for *D. melanogaster* (Zheng et al., 2018; Dorkenwald et al., 2022) and a connectome for its ventral nervous system (Azevedo et al., 2022).

However, EM data does not directly give us information about gene expression and as a result key neuronal features such as transmitter identity. The action a neuron has on its downstream targets, namely excitation or inhibition, depends on the transmitter it releases, yet transmitter identity is unknown for a majority of neuronal cell types in the connectomes of *D. melanogaster*. It is expected that each neuron expresses only one transmitter (*Dale’s law*) (Eccles, 1976; Dale, 1934), ocasionally a small selection of transmitters. In addition, neurons of the same neuronal cell type will express the same transmitter between different animals, *i*.*e*., expression is stereotyped (Bates et al., 2019). Before release, transmitters are packaged into different types of vesicles at synaptic sites. The so-called *classical* fast-acting transmitters (acetylcholine, glutamate, and GABA) are the most common and are contained in small, clear vesicles. Monoamines such as dopamine, norepinephrine, octopamine and serotonin are packaged into pleomorphic clear core or small dense core vesicles (Goyal and Chaudhury, 2013). The many neuropeptides that *D. melanogaster* express, such as cholecystokinin, galanin, neurokinin, neuropeptide F and oxytocin, are contained in larger dense core vesicles.

In mammals and some large invertebrates it is sometimes possible for humans to distinguish between different clear core and dense core vesicles and thus identify the transmitter of a given synaptic site. Among clear core vesicle, human experts can distinguish excitatory and inhibitory transmitters by how elliptical their synaptic vesicles are (Tao et al., 2018; Atwood et al., 1972; Uchizono, 1965), though they have not yet to our knowledge been automatically predicted at scale in these species.

In small invertebrates, such as *D. melanogaster*, it is so far unknown whether synaptic phenotype is sufficient to consistently determine transmitter identity, especially different varieties of clear core vesicles. Synapse transmitter identity cannot be consistently identified by human annotators from EM alone.

As a result, adding transmitter identities to connectomic data requires molecular biology and light microscopy pipelines, in which gene transcripts or proteins involved in the pathway of interest have been made visible using fluorescent probes (Hyatt and Wise, 2001; Long et al., 2017; Meissner et al., 2019), or by RNA sequencing (Henry et al., 2012; Konstantinides et al., 2015; Davie et al., 2018; Davis et al., 2020). However, this approach is very difficult to scale to an entire nervous system and conventional means of profiling many cells cannot be used. Fundamentally, this is because the majority of morphological neuronal types cannot yet be linked to RNA expression identities (Bates et al., 2019), though they can be unambiguously linked to connectomic reconstructions (Bates et al., 2020b).

The fly nervous system is staggeringly diverse. For *D. melanogaster*, we want to know the transmitter identity of each of ∼115,000 neurons in the brain alone (Mu et al., 2021) and ∼32,500 in the central brain (i.e. discounting the purely optic regions, Schlegel et al. in prep). The central brain’s ∼32,500 neurons can be decomposed into perhaps ∼6,000 isomorphic neuronal cell types (Bates et al., 2019), each of which could have a different transmitter expression from its neighbours and each plays a different and unique role in the brain’s circuits. Of these, the expression of only ∼400 cell types are known after a few decades of work. To discover the transmitter identity of further cell types, investigators require transmitter expression data that can be unambiguously linked to single neuron morphologies. To that end, well-characterized, sparse genetic driver lines that only target a few neurons must be investigated, one at a time, followed by accurate morphological matching to EM tracings using neuron similarity tools, such as NBLAST (Costa et al., 2016) or color MIP searches (Otsuna et al., 2018). It is an arduous pipeline that has not been reliably automated.

Here, we show that it is possible instead to determine the primary transmitter of a given neuron in the *D. melanogaster* brain from the phenotype of its synaptic sites in EM alone. We learn a mapping *f*: *x* → *y*, where *x* is a local 640 × 640 × 640nm^3^ 3D EM volume with a synaptic site at the center and *y* the neurotransmitter of the corresponding neuron (see Fig. 1). To learn this mapping, we generated training data set of pairs (*x, y*). This involved matching of neuron visualisations from light microscopy to EM neuronal reconstructions, for which we had densely annotated presynaptic sites. We trained a neural network to classify each synapse into one of the following six transmitters: GABA, acetylcholine, glutamate, serotonin, octopamine, and dopamine. We deploy this method on two different data sets, the full adult fly brain (FAFB) (Zheng et al., 2018) and the HemiBrain (Scheffer et al., 2020a). In both data sets, we find that this method is able to classify the transmitter of any given synapse with high accuracy (88% for FAFB and 79% for HemiBrain, see Fig. 2 for details). We have also generated results for a third data set (MaleVNC), a full adult male ventral nervous system being reconstructed at Janelia Research Campus, where our results are consistent with Lacin et al. (2019).

**Figure 1:**
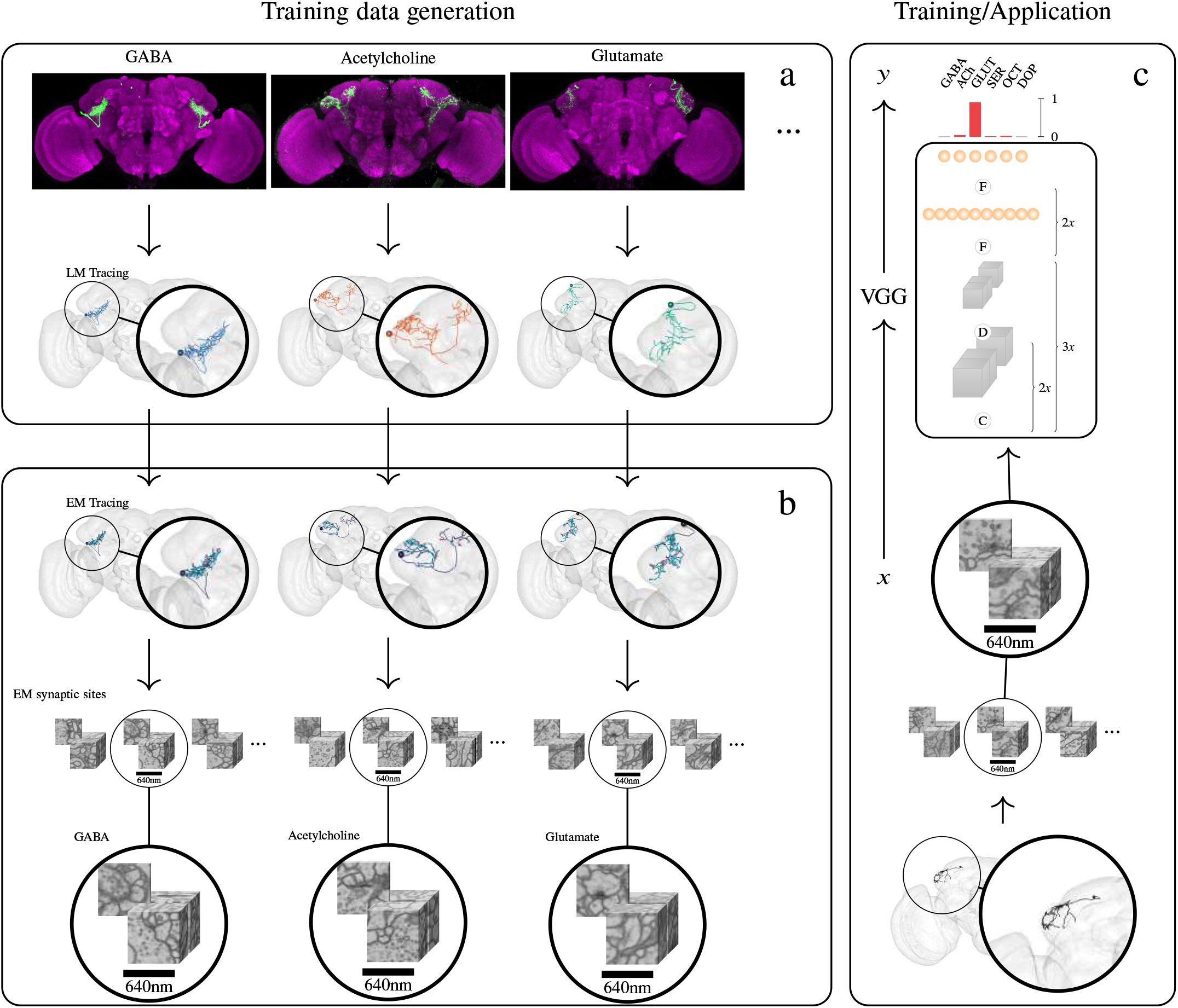
Method Overview. We assemble a data set of Catmaid neurons with known transmitter in the *D. melanogaster* whole brain EM data set (FAFB) from the literature and retrieve corresponding synaptic locations from the subset of skeletons that have been annotated in the Catmaid collaboration database1. Typically, neurons are genetically tagged to identify their transmitter identity and reconstruct their coarse morphology using light microscopy (**a**). Light microscopy tracings of neurons are then matched to corresponding neuron traces in Catmaid, and synaptic locations are annotated, resulting in a data set of EM volumes of synaptic sites with known transmitter identity (**b**). We use the resulting pair (*x, y*), with *x* a 3D EM volume of a synaptic site and *y* ∈ {GABA, acetylcholine, glutamate, serotonin, octopamine}, dopamine, the transmitter of that synaptic site, to train a 3D VGG-style deep neural network to assign a given synaptic site *x* to one of the six considered transmitters. We use the trained network to predict the transmitter identity of synapses from neurons with so far unknown transmitter identity in the *D. melanogaster* Catmaid data set (**c**). C, D, and F denote convolution, down-sampling, and fully connected layers respectively. We used the same method with the HemiBrain data set

**Figure 2:**
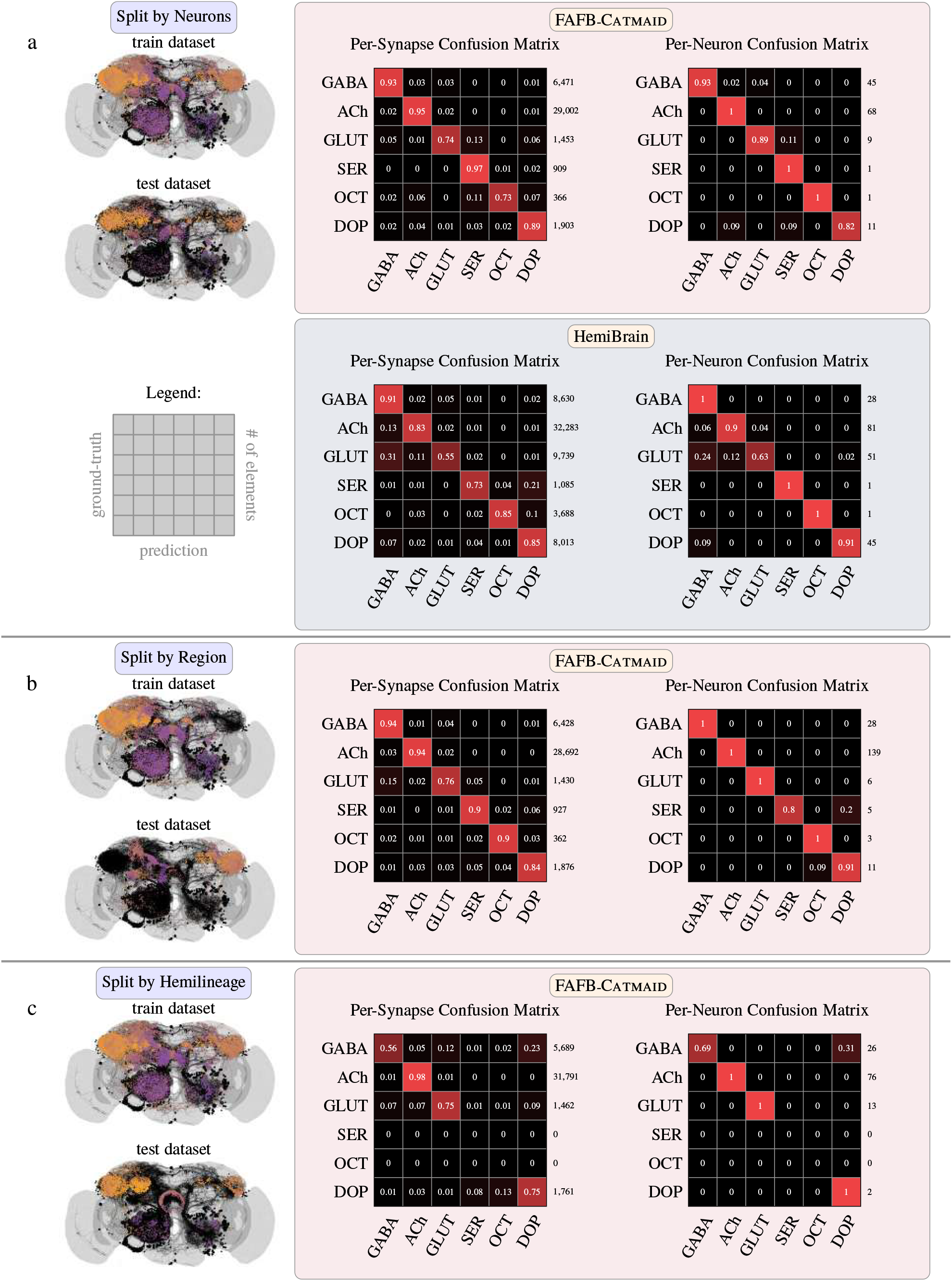
Illustration of the train and test data sets and the accuracy of the trained classifier on a per-synapse and per-neuron basis. **(a)**: Left: visualization of the train and testing data (split by entire neurons) used for the main results in this manuscript. Synapse locations are color coded according to their z-depth (perpendicular to viewing plane). Right: confusion matrices of the trained classifier on the testing data, shown per synapse and as a majority vote per neuron, on data sets FAFB and HemiBrain. In order to be able to have a meaningful majority vote we only consider neurons with more than 30 synapses for the per-neuron confusion matrices. **(b)**: Classification results on an alternative train and testing data (split by brain regions) on FAFB to investigate potential brain region confounds. **(c)**: Same as (b), but split by hemilineage. For the hemilineage split it was not possible to generate a fully balanced split and as a result there are no serotonin and octopamine neurons in the test set, as indicated by the zero-filled rows.

Our predictions allowed us to ask a set of biological questions to assess the inventory of the *D. melanogaster* central brain connectomes with regard to transmitter expression. We make use of tens of thousands of neurons semi-automatically reconstructed in the HemiBrain and from the ongoing semi-automatic reconstruction projects in FAFB (via flywire.ai, Dorkenwald et al. (2022)) and MaleVNC. We find that neurons that express different transmitters can differ in key cellular features as well as their connectivity within a neural system. We also find that transmitter usage is consistent between axon and dendrite, members of the same neuronal cell type, sides of the brain, and identifiable neurons between two different flies. In addition, we find that the principle that a single fast-acting transmitter is expressed per developmental unit (*Lacin’s law*, Lacin et al. (2019)) largely holds true in the *D. melanogaster* central brain, with a small number of interesting deviants. We further find that neuron populations from the main transmitter groups differ in their polarisation, projection, mitochondria count, excitation-inhibition balance, and input profiles.

We have made transmitter predictions for FAFB and Hemi-Brain publicly available to help accelerate connectome interpretation in *D. melanogaster*.

## 2 Results

### 2.1 Assembling Ground Truth Presynapse Data for *D. melanogaster*

We built our ground-truth data from manually placed presynaptic sites in FAFB using the annotation software CATMAID by the Catmaid-community (Zheng et al., 2018; Schneider-Mizell et al., 2016) (see Acknowledgements). We then make predictions across automatically predicted presynapses in FAFB (Buhmann et al., 2019) and conduct biological analyses from neurons semiautomatically built in FlyWire by the flywire.ai community (see Acknowledgements) because ten-fold fewer reconstructions are available in Catmaid. FlyWire is an active project that aims to complete the first full brain adult insect connectome (Dorkenwald et al., 2022). In addition, we used automatically detected presynapses and semi-automatically reconstructed neurons in a second data set, HemiBrain (Scheffer et al., 2020a).

In order to begin learning the transmitter identity for presynapses we collected many thousands of synaptic locations with known transmitter identity. To that end, we created a list of 356 neuronal cell types from 21 studies (see supplemental data file 7.1.1) and identified as many of them as we could in the Catmaid and HemiBrain data sets (Dolan et al., 2019; Aso et al., 2014; Palavicino-Maggio et al., 2019; Davis et al., 2020; Henry et al., 2012; Tanaka et al., 2012; Nässel and Elekes, 1992; Wolff et al., 2015; Mao and Davis, 2009; Niens et al., 2017; Liu et al., 2009; Turner-Evans et al., 2020; Guo et al., 2016; Lu et al., 2022; Frenkel et al., 2017; Shafer et al., 2006; Senapati et al., 2019; Krashes et al., 2009; Busch et al., 2009; Liu et al., 2019; Dacks et al., 2006; Haynes et al., 2015; Hulse et al., 2021; Scheffer et al., 2020a) (see Acknowledgements).

The transmitter expression evidence in these studies comes from detecting gene expression related to transmitter usage (minority) or immunohistochemistry (majority) experiments. For these studies, neurons were picked (transcriptomics) or stained (immunohistochemistry) with guidance from green fluorescent protein (GFP) expression in a GAL4/split-GAL4 line (Jenett et al., 2012). These genetic driver lines target transgene expression to a small constellation of discriminable cell types, or even individual cell types or neurons. Typically, brains are dissected out and incubated with primary antibodies (*e*.*g*., anti-VGlut, anti-GABA, anti-ChAT), followed by secondary antibodies which have a fluorescent tag to visualise the primary antibody, or else native RNA transcripts are targeted by a fluorescent probe (*in situ* hybridization). The proteins/transcripts related to certain transmitter expressions are thus labelled across the brain. If they co-localize with the GFP signal for the GAL4/split-GAL4 line of interest, those neurons are considered to express that transmitter (a commonly used, full step-by-step protocol can be found at https://www.janelia.org/project-team/flylight/protocols). Note that individual studies usually only tested single transmitters and did not show negative staining. As a result, there is limited data for co-transmission of multiple transmitters in a single neuron and we therefore assume Dale’s law, *i*.*e*., no co-transmission of transmitters within a single neuron. Other methods involve RNA sequencing and include TAPIN-seq (Davis et al., 2020), and in these cases it is more possible to accumulate data indicative of transmitter co-release.

Many neurons break from Dale’s law, which we can see in single cell sequencing data (Croset et al., 2018), but for the vast majority of data points in these transcriptomic data we do not know the neuron’s morphology and therefore can not link them to connectomic reconstructions. However, there are only a small number of identified cell types—for which we know both transmitter expression and neuronal morphology—in which co-transmission is known to occur, *e*.*g*., Kenyon cells (acetylcholine/sNPF/others) (Croset et al., 2018), DPM (GABA/serotonin/amnesiac) (Waddell et al., 2000), Mi15 (acetylcholine/serotonin) (Davis et al., 2020) and PPL101-6 DANs (dopamine/serotonin) (Mao and Davis, 2009). We excluded those cell types from our analysis. While all octopaminergic neurons are known to co-transmit glutamate (Sherer et al., 2020; Croset et al., 2018), we still treat them as their own octopamine category.

The HemiBrain data set (https://neuprint.janelia.org/?dataset=hemibrain:v1.2.1) had already been cell typed and largely linked to names in use in the literature (Scheffer et al., 2020a). Each reconstruction has associated automatically detected presynapses (Huang et al., 2018). In Catmaid(https://neuropil.janelia.org/tracing/fafb) we already knew many of the olfactory, thermosensory, mushroom body and some central complex cell types (Bates et al., 2020b; Dolan et al., 2018; Felsenberg et al., 2018a; Frechter et al., 2019; Huoviala et al., 2018; Dolan et al., 2019; Marin et al., 2020; Sayin et al., 2019; Turner-Evans et al., 2019; Zheng et al., 2018) and sought to identify others in the visual system and elsewhere using matches to light microscopy and leveraging expert knowledge in the research community (see Acknowledgements). In these studies, the authors had already linked some of their reconstructed cell types to immunohistochemical data (Aso et al., 2014; Bräcker et al., 2013; Busch et al., 2009; Davis et al., 2020; Dolan et al., 2019; Ito et al., 2013; Lai et al., 2008; Okada et al., 2009; Shinomiya et al., 2015; Tanaka et al., 2012; Wilson and Laurent, 2005b). Instead of using the FlyWire platform (https://ngl.flywire.ai/), its automatic segmentation of FAFB (Dorkenwald et al., 2022) and its synapses (Buhmann et al., 2019) we used manually created Catmaid reconstructions with manually annotated presynapses from Catmaid (http://www.catmaid.org) (Saalfeld et al., 2009; Schneider-Mizell et al., 2016). We did this to ensure that we used a large number of high fidelity synapses. We matched 3025 Catmaid neuronal reconstructions to cell types with a known transmitter. We also matched 5902 HemiBrain reconstructions to cell types with a known transmitter. For Catmaid neurons, we increased the number of labelled synapses by furthering manual synaptic assignments on the neurons of interest. Synapses were annotated at presynaptic sites, defined by T-bars, vesicles and a thick dark active zone by a synaptic cleft (Prokop and Meinertzhagen, 2006). We scored each continuous synaptic cleft as a single presynapse regardless of its size or the number of associated T-bars.

In total, the assembled Catmaid ground truth data set contains 153,593 acetylcholine presynapses (587 neurons), 7,953 glutamate presynapses (50 neurons), 32,836 GABA presynapses (175 neurons), 9,526 dopamine presynapses (83 neurons), 4,732 serotonin presynapses (5 neurons), and 2,924 octopamine presynapses (6 neurons) (see supplemental data file 7.1.2). The assembled Hemi-Brain ground truth data set was larger, thanks to the available cell type annotations, more extant reconstructions and our shift to use automatically detected presynapses. It contains 451,033 acetylcholine presynapses (3,094 neurons), 75,239 glutamate presynapses (218 neurons), 80,732 GABA presynapses (242 neurons), 117,054 dopamine presynapses (310 neurons), 70,460 serotonin presynapses (38 neurons), and 46,017 octopamine presynapses (21 neurons).

Due to a relative paucity of presynapses we did not use our annotations for the other transmitters we had collated, namely: Allatostatin-a (2 neurons in the HemiBrain), Corazonin (6), Drosulfakinin (6), glycine (9), insulin (23), IPNa (2), NPF (3), and SIFamide (4). Histamine is also used in the *D. melanogaster* brain, primarily by photoreceptor neurons. More transmitters than these are also used but the exact number of transmitters and the number of neurons that express them instead of one of the six for which we had sufficient ground truth data, is unknown.

### 2.2 Train and Test data sets

For each transmitter y ∈ *{*GABA, acetylcholine, glutamate, serotonin, octopamine, dopamine}, we divide the data into test, train, and validation set by randomly assigning entire neurons, each containing multiple synapses. We refer to this split as *neuron split* in the following. We use 70% of neurons for training, 10% for validation and the remaining 20% for testing. Splitting the data set by entire neurons, instead of randomly sampling pre-synapses, mirrors the real world use case in which we typically know the transmitter of an entire neuron and are interested in the transmitter of a different neuron.

In order to test how well the proposed methods generalizes across morphologically distinct cells and regions, and to exclude potential confounds, we also generated two additional splits that separate the data by hemilineage (*hemilineage split, i*.*e*., neurons grouped by shared developmental origin) and brain region (*brain region split, i*.*e*., into standard neuropils (Ito et al., 2014)) respectively. To this end we found the optimal split between entire hemilineages and brain regions, such that the fraction of synapses for every transmitter in the train set is close to 80% of all synapses of that transmitter. We further used randomly selected 12.5% of the training synapses (10% of the entire data set) for validation.

### 2.3 Network Architecture and Training

We used a 3D VGG-style (Simonyan and Zisserman, 2014) network to predict the transmitter identity from a 3D EM input cube of edge length 640 nm with a synaptic site at its center. The network consists of four functional blocks, each consisting of two 3D convolution operations, batch normalization, ReLU non-linearities and subsequent max pooling with a downsample factor of 2, except for FAFB where we limit downsampling to the x and y dimensions for the first three blocks to account for anisotropy. The last block is followed by three fully connected layers with dropout (p=0.5) applied to the last one. We trained the network to minimize cross-entropy loss over the six classes (GABA, acetylcholine, glutamate, serotonin, octopamine and dopamine), using the Adam optimizer (Kingma and Ba, 2014). We trained for a total of 500,000 iterations in batches of size eight and select the iteration with highest validation accuracy for testing. For an illustration of the used network architecture see Fig. 1.

### 2.4 Accuracy of Neurotransmitter Predictions

We tested the classifier on our held-out test sets. For the *neuron split*, the test set consists of a total of 40,104 synapses from 185 neurons that the network was not trained on. We achieved an average per-transmitter accuracy of 87% for the transmitter prediction of individual synapses on FAFB and 78% on HemiBrain. Since we assume that each neuron expresses the same neurotransmitters at its synaptic sites, we can quantify the per neuron accuracy of the presented method. To this end, we assigned each neuron with more than 30 synapses in the test set a transmitter by a majority vote of its synapses, leading to an average accuracy of 94% for the transmitter prediction per neuron on FAFB and 91% on HemiBrain. For the *hemilineage split*, we find an accuracy of 75% for individual synapses and 92% for entire neurons. The *Brain Region split* evaluates to 88% synapse classification accuracy and 95% neuron classification accuracy. A per-class overview can be seen in Fig. 2, for a summary of the results and data splits see Table 1.

**Table 1:** Overview of the three data splits used for evaluation of the classifier. Shown are the number of synapses for training, testing and validation as well as average synapse and neuron classification accuracy on the test set for each data split.

|  | Neuron split | Hemilineage split | Brain Region split |
| --- | --- | --- | --- |
| Train | 140,565 | 140,868 | 138,982 |
| Test | 40,104 | 40,703 | 39,715 |
| Validation | 20,084 | 19,182 | 19,858 |
| Avg. Synapse Accuracy | 87% | 75% | 88% |
| Avg. Neuron Accuracy | 94% | 92% | 95% |

### 2.5 Classifier Feature Analysis

The high classifier accuracy on the held-out testing data sets is surprising, since human annotators can not reliably identify the transmitter identity of synapses from their EM images alone. Although coarse classifications can be done by eye (*e*.*g*., clear core vesicles are more likely to be associated with classical transmitters than dense core vesicles) (Thureson-Klein, 1983; Kemali, 1976), an explanation for the classifier’s ability to discriminate between all six transmitters is lacking.

We, therefore, sought to visualize which features the classifier uses to discriminate between different transmitters. The motivation for identifying those features is two-fold: First, we can not rule out that the classifier leveraged confounds that are potentially present in our training and testing data. It is, for example, conceivable that different brain regions have slightly different visual properties and correlate with specific transmitters more than others. We were able to rule this particular confound out (see Fig. 2b), but similar biases could in principle be present in our training and testing data. Second, a feature visualization can shed light on the interplay of structure and function by identifying sub-cellular structures that are predictive of certain transmitters.

To analyze the features the classifier makes use of, we are particularly interested in so-called *post-hoc* attribution methods, *i*.*e*., methods that try to explain decisions of an already trained classifier. Several methods have been proposed for post-hoc attribution on images, *e*.*g*., Input*Gradients, IntegratedGradients, DeepLift, or GradCAM (Shrikumar et al., 2016; Sundararajan et al., 2017; Ancona et al., 2018; Selvaraju et al., 2017). Those methods aim to derive an *attribution map* for a single input image, such that areas of the image that were crucial for making a classification decision are highlighted. We will refer to those methods as *single-input* attribution methods.

We found that the attribution maps generated by those methods on EM images of synapses could not clearly be interpreted. The areas highlighted by the attribution maps vary greatly between different methods and are generally too large to point to individual sub-cellular structures like vesicles (data not shown). We hypothesize that this is mostly because single-input attribution methods are subject to highlighting so-called “distractors”, *i*.*e*., non-class-specific features that are shared between images of different classes (*e*.*g*., features stemming from different orientations of the synapse, section thickness, or intensity variations) (Kindermans et al., 2017).

We therefore developed a novel attribution method to highlight *differences* between images of two distinct classes by focusing solely on class-relevant features and disregarding distractors (Eckstein et al., 2023): Given a real image ***x***_R_ of a class *y*_R_, we first create a *counterfactual* image ***x***_C_ by translating ***x***_R_ into an image of another class *y*_C_ using a CycleGAN (Zhu et al., 2017), resulting in *paired* images of different neurotransmitter types. Crucially, this domain translation of an image from class *y*_R_ to class *y*_C_ keeps class-irrelevant distractors (like the orientation of the synapse in the image) intact. Class-relevant features, however, are changed due to the adversarially trained discriminator of the CycleGAN. We use our previously trained classifier to confirm that the domain translation was successful: we filter all images such that ***x***_R_ is classified as *y*_R_ and the counterfactual ***x***_C_ as *y*_C_. We then identify a small region in ***x***_C_, such that swapping its content with the one from ***x***_R_ also changes the prediction of the trained classifier (see examples in Fig. 3B). To this end, we search for a minimal binary mask ***m***, such that the hybrid image ***x***_H_ = ***m*** · ***x***_R_ + (1 – ***m***)· ***x***_C_ is classified as *y*_R_. To find ***m***, we use a modified version of the DeepLift method, where we use the counterfactual ***x***_C_ as the “neutral” reference (Eckstein et al., 2023).

**Figure 3:**
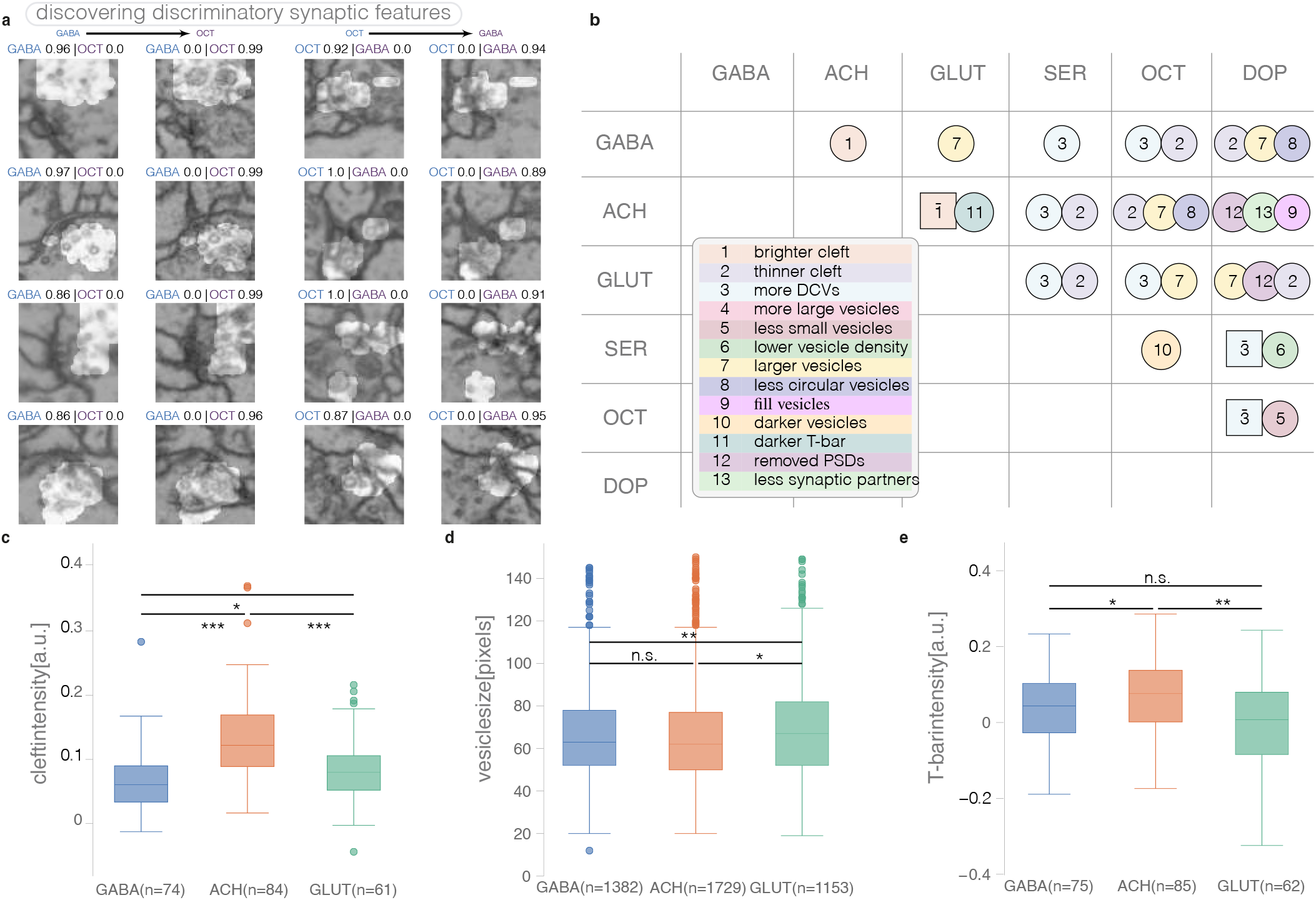
Classifier feature analysis using a discriminative attribution method. **(a)**: Example translations of real synapse images (blue label) to fake counterfactual images (purple label). Attribution masks indicating the most important changes between the two classes are shown as highlights. Classifier scores are shown above each image for the real and counterfactual class. Left two columns show the translation of real GABA synapses into counterfactual octopamine synapses, right two columns the same for octopamine to GABA. **(b)**: Pairwise differences between transmitters, found through manual inspection of real and counterfactual images as shown in (a). Squares indicate the opposite of the features listed in the legend. **(c)**: Quantitative comparison of the cleft intensity between transmitters GABA, acetylcholine, and glutamate on the original synapse images, confirming the findings in (b). **(d)** and **(e)**: Same as (c) for vesicle sizes and t-bar intensities. (*n*.*s*.: not significant, *: *p* ≤ 0.05, **: *p* ≤ 0.001, ***: *p* ≤ 0.0001,)

This method allowed us to manually identify at least one distinguishing feature between each pair of transmitters (see Fig. 3A). We only included features that we observed consistently in both directions, *e*.*g*., translating a real GABA image to acetylcholine results in a brighter cleft and translating from a real acetylcholine image to GABA in a darker cleft. Being able to compare two neurotransmitter identities in paired images also allowed us to observe features as subtle as sub-pixel changes in vesicle diameters (*e*.*g*., between GABA and glutamate), which would be near impossible to pick up in unpaired images.

We confirmed the identified features between the classical transmitters GABA, acetylcholine, and glutamate on the original synapse images. To this end, we manually segmented the synaptic cleft, vesicles, and t-bars of 222 synapse images (75 GABA, 85 acetylcholine, 62 glutamate; annotators were blind to the neurotransmitter type). We did indeed find strong support for each of the identified features between those transmitters (Fig. 3C, D, and E): acetylcholine has a brighter cleft than GABA and glutamate (*p* ≤ 0.0001), glutamate has larger vesicles than GABA (*p* ≤ 0.001), and glutamate has a darker t-bar than acetylcholine (*p* ≤ 0.001).

### 2.6 Comparison of Predictions for Homologs Across Data Sets

The neuronal inventory of the insect brain can be broken down into isomorphic cell types (Fig. 5d), each of which usually contains only a few neurons (Bates et al., 2019) (mean, 2.3, s.d., 2.6 per hemisphere per cell type for the hemibrain data set, excluding the largest outlier classes: Kenyon cells, sensory receptor neurons, ER neurons and EPG neurons in the HemiBrain data set). These cell types are duplicated on both hemispheres. In many cases, cell types comprise only a single neuron (41% of the hemibrain data set). We refer to these neurons as *singletons*. Because these singletons have a unique morphology between both sides of the brain, they can be matched unambiguously to their homolog on the other side of the same brain, or a homolog in the brain of another member of its species (Fig. 4a). In both cases, the singleton will have developed from a homolog neuroblast during embryonic or larval development and therefore have a similar gene expression profile (Yang et al., 2016), meaning that we strongly expect matched singletons to each express the same transmitter - both between hemispheres of the same brain (testable within the FlyWire data set) and between two different brains (testable by comparison to the HemiBrain data set). For this reason, using matched singletons across brain hemispheres provides a natural mechanism by which to test whether our transmitter predictions are consistent.

**Figure 4:**
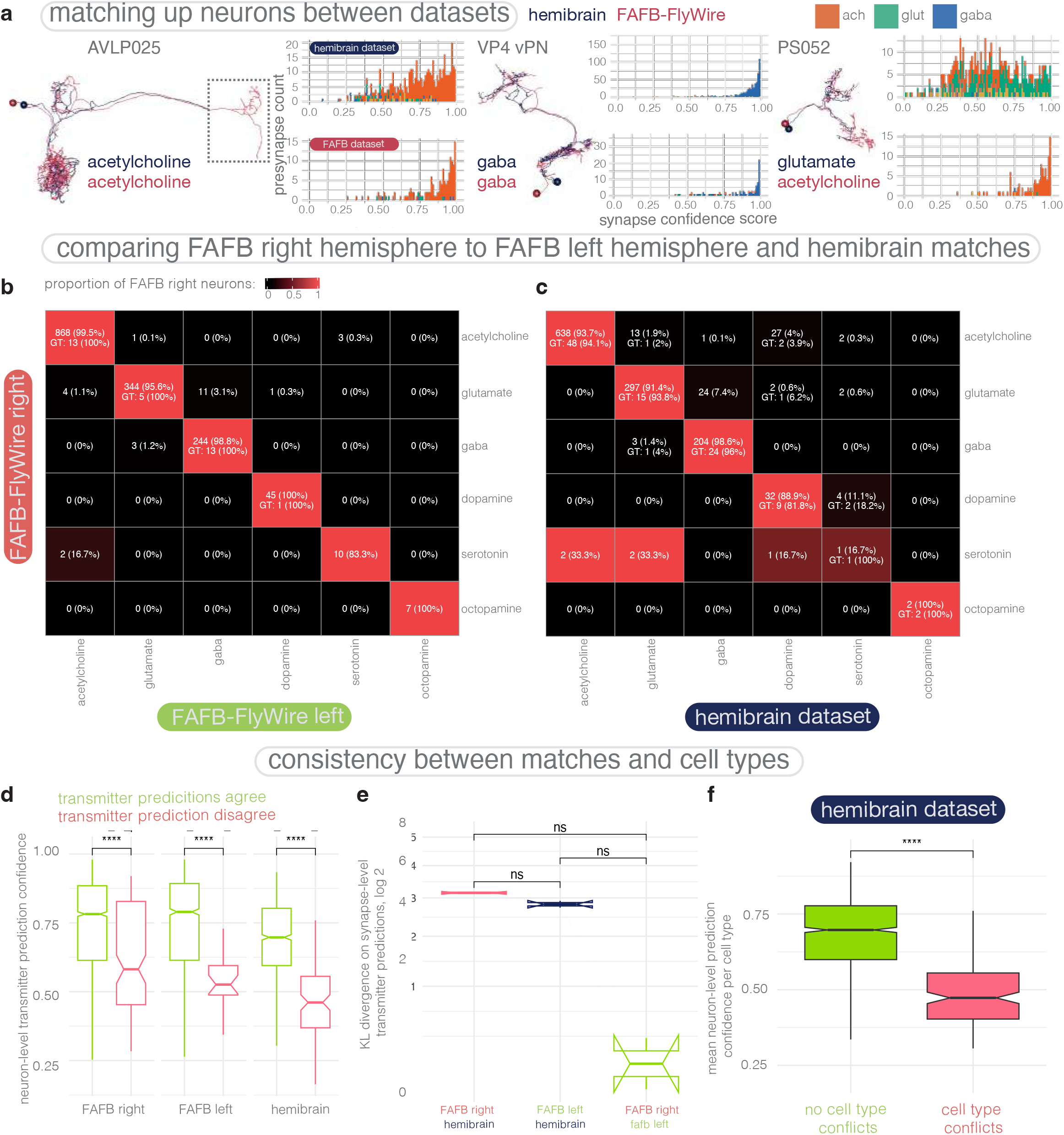
Comparing neurons’ transmitter usage predictions between connectome data sets from separate animals (FAFB-FlyWire and HemiBrain) and between two hemispheres in the same data set (FAFB-FlyWire). **(a)** Neurons can be matched up between FAFB-FlyWire and the HemiBrain convincingly. Images of co-registered, matched neurons between the HemiBrain data set (orange) and the FAFB-FlyWire data set (blue). These neurons belong to the same cell type and hemisphere, in both connectome data sets. Histograms show synapse-level transmitter prediction scores for both neurons in each of our exemplar matched pairs. The cell type AVLP025 can be matched despite missing data in the HemiBrain connectome (left, grey dashed box). The cell type PS052 has conflicting neuron-level transmitter predictions between FAFB-FlyWire (acetylcholine) and HemiBrain (glutamate) despite a strong morphological match. **(b)** FAFB-FlyWire right hemisphere to FAFB-FlyWire left hemisphere matches, 1,543 pairs. Copies of the same neuron on the right and left sides of the fly brain have been matched, and their neuron-level transmitter predictions are compared. Left, a confusion matrix comparing FAFB-FlyWire right and FAFB-FlyWire left neuron-level transmitter predictions. Cells coloured by the proportion of FAFB-FlyWire right neurons of each transmitter type, that are matched to a FAFB-FlyWire left homolog of each transmitter type (row normalised). There are 878 acetylcholine, 382 glutamate, 272 GABA, 45 dopamine, 18, serotonin, 7 octopamine matches by the FAFB-FlyWire right side prediction, of which 40 (2.5%) disagree with the FAFB-FlyWire left side prediction. **(c)** FAFB-FlyWire right hemisphere to HemiBrain right hemisphere matches, 1,257 pairs. Copies of the same neuron on the same hemisphere across the FAFB-FlyWire (rows) and HemiBrain (columns) data sets, and their neuron-level transmitter predictions are compared. Left, a confusion matrix comparing FAFB-FlyWire right and HemiBrain right neuron-level transmitter predictions. Cells coloured by the proportion of FAFB-FlyWire right neurons of each transmitter type, that are matched to a HemiBrain homolog of each transmitter type (row normalised). There are 630 acetylcholine 329 glutamate, 209 GABA, 39 dopamine, 7, serotonin, 4 octopamine matches by the FAFB-FlyWire right side prediction, of which 94 (7.7%) disagree with the HemiBrain prediction. Cells indicate number of neurons that were also represented in our ground truth synapse training data, albeit for FAFB-FlyWire neurons these synapses would have been different (we used manually placed presynapse markers for FAFB training, here we examine automatically detected presynaptic sites (Buhmann et al., 2019)). **(d)** neuron-level transmitter prediction scores between matched neurons that have (red, right) or do not have (green, left) a conflict between their neuron-level transmitter predictions, across all three hemispheres. Number of matches/mismatches across all comparisons for FAFB-FlyWire right, 2650:170 FAFB-FlyWire left, 1562:40, and HemiBrain neurons, 1088:130. Data compared using Wilcoxon two-sample tests. Significance values: ns: p > 0.05; *: p <=0.05; **: p <= 0.01; ***: p <=0.001; ****: p <= 0.0001. **(e)** Comparison of similarity scores (Kullback–Leibler divergence) for matches based on their synapse-level transmitter prediction scores. Red, FAFB-right-HemiBrain matches, blue, FAFB-left-HemiBrain matches, green, FAFB-right-FAFB-left matches. The higher the score, the less similar the probability distribution of synapse-level transmitter prediction scores (where for each synapse there is a confidence score associated with each of our 6 possible transmitter predictions). Y axis is log2 transformed. **(f)** The neuron-level transmitter prediction consistency among HemiBrain neuronal cell types. Plot is for cell types with more than one member on the right (*i*.*e*., the only full) hemisphere in the HemiBrain data set. Plot is for cell types with more than one member on the right (*i*.*e*., the only full) hemisphere in the HemiBrain data set. Green, the mean neuron-level transmitter prediction confidence for cell types where there is no conflict, *i*.*e*., all members of the type are predicted to use the same transmitter (N=1,091). Red, the mean neuron-level transmitter prediction confidence for cell types where there is a conflict, *i*.*e*., not all members of the type are predicted to use the same transmitter (N=95). Box plots show the median (horizontal line), the lower and upper hinges correspond to the first and third quartiles (the 25^th^ and 75^th^ percentiles). Whiskers extend to 1.5 times the inter-quartile range (IQR). The notch displays a confidence interval around the median. This interval is 1.58*IQR/sqrt(n). If the notches do not overlap, the medians significantly differ. FAFB data built from FlyWire reconstructions.

**Figure 5:**
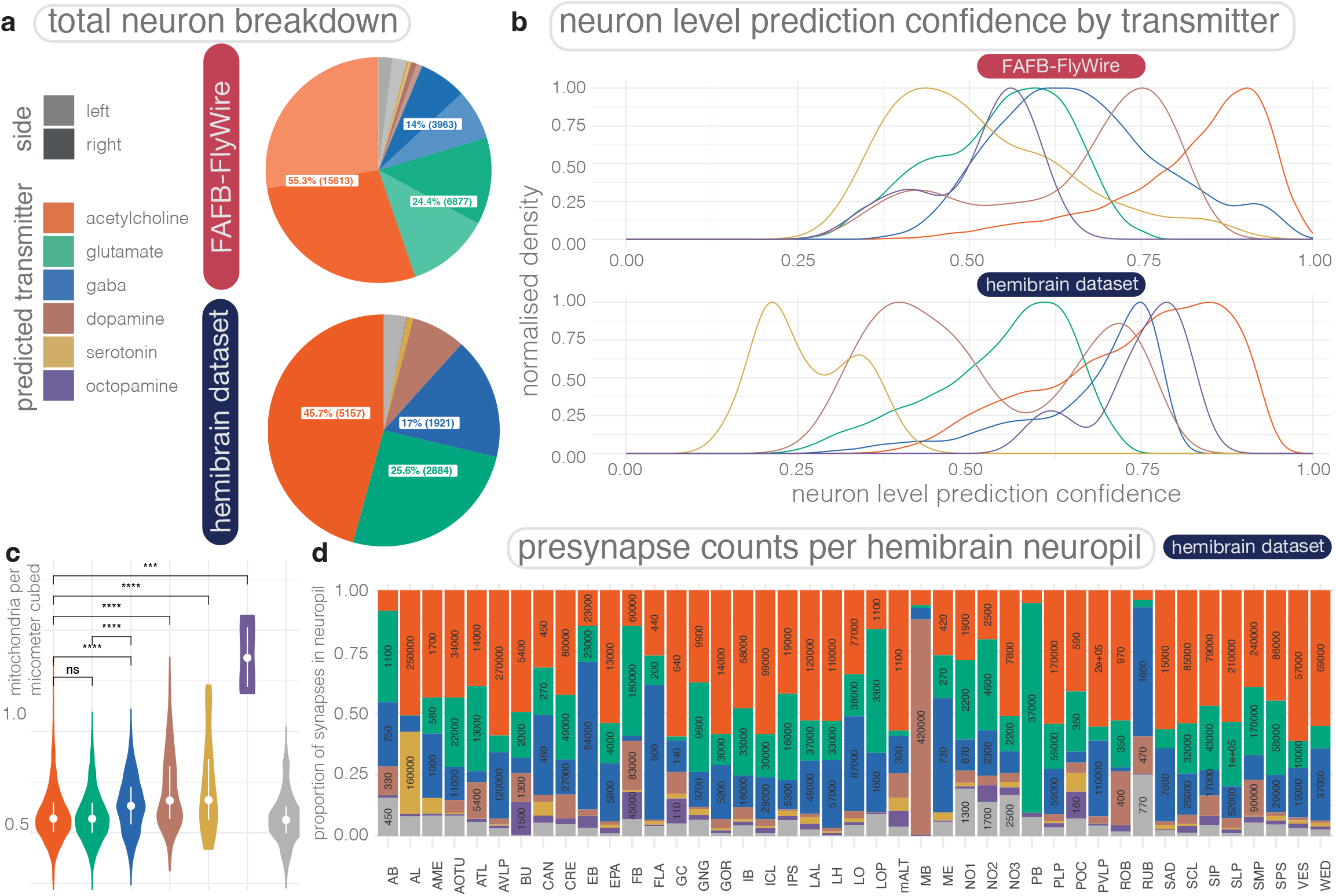
Neuronal transmitter expression across the HemiBrain and FAFB-FlyWire partial connectomes. **(a)** A breakdown of neuron-level transmitter expression among the 11,277 neurons of the HemiBrain (upper) and 27,706 well reconstructed neurons across FAFB-FlyWire (lower). FAFB-FlyWire reconstructions were determined to be well reconstructed if a human annotator could confirm they appeared to have a full dendrite, axons, cell body fibre tract and cell body (soma), and the neuron contained at least 100 presynapses. In the HemiBrain connectome there are 24,709, which we filtered down to 11,277 by excluding neurons without the HemiBrain project status label “Traced”, neurons with less than 100 presynapses and removing neurons with a large part of their arbor cut from the data set, including neurons with the terms: LP, LC, LT, DN and LLP in their cell type label. We excluded Kenyon cells from both data sets. The Kenyon Cell lineages (MBp1-4) are indicated by a red bar, these lineages are likely cholinergic rather than dopaminergic (see main text). **(b)** Scaled density plots (max-normlised, smoothed histograms) for the distribution of neuron-level transmitter prediction scores across our different transmitter types. The Y axis gives a max normalised density of neuron number. **(c)** Mitochondria density neuron-level transmitter predictions. Violin plots show the number of automatically detected mitochondia (Plaza et al., 2022) per micron cubed. Volume measures per neuron originate from the HemiBrain’s automatically reconstructed 3D neuron volumes (Scheffer et al., 2020a). A mitochondria detection is currently only available in the HemiBrain data set. The mean number of mitochondria per neuron is 245, s.d. 275. **(d)** Breakdown of neuron-level transmitter predictions by brain region. Plot shows the proportion of synapses in each HemiBrain neuropil that belong to a neuron of a given neuron-level transmitter prediction (colors). A total of ∼4,000,000 were assigned a neuropil and neuron-level transmitter prediction, which helps buffer erroneous synapse-level transmitter predictions. Number labels give the total number of synapses in each group. Not all the standard neuropils (Ito et al., 2014) are shown because the HemiBrain only comprises ∼1/3 of the central brain. FAFB neuropils not shown because we only used 27,706 of an estimated 32,000 FAFB-FlyWire central brain neurons (Croset et al., 2018), and therefore could not be sure we have a representative sample of synapses from each neuropil. Kenyon cells of the mushroom body have been excluded from this analysis. Only synapses predicted to be located on an axon or dendrite are included. Note that all known octopaminergic neurons in the fly brain are also glutamatergic (Sherer et al., 2020), so the octopamine label actually indicates octopamine and glutamate co-transmission. Neuropils (Ito et al., 2014): AB, asymmetric body, AL, antennal lobe, AME, acessory medulla, AOTU, anterior optic tubercle, ATL, antler, AVLP, anterior ventrolaterla protocerebrum (incomplete), BU, bulb, CAN, cantle, CRE, crepine, EB, ellipsoid body, EPA, epaulette, FB, fan-shaped body, FLA, flange, GC, great commissure (incomplete), GNG, gnathal ganglion (incomplete), GOR, gorget, IB, inframedial bridge, ICL, inferior clamp, IPS, inferior posterior slope, LAL, lateral acessory lobe, LH, lateral horn, LO, lobula (incomplete), LOP, lobula plate (incomplete), ME, medulla (incomplete), NO1, nodulus compartment 1, NO2, nodulus compartment 2, NO3, nodulus compartment 3, PB, protocerebral bridge, PLP, posterior lateral protocerebrum, POC, posterior optic commissure, PVLP, posterior ventrolateral protocerebrum (incomplete), ROB, round body, RUB, rubus, SAD, saddle, SCL, superior clamp, SIP, superior intermediate protocerebrum, SLP, superior lateral protocerebrum, SMP, superior medial protocerebrum, SPS, superior posterior slope, VES, vest, WED, wedge. Violin plots show the median (dot), the first and third quartiles (lines, the 25^th^ and 75^th^ percentiles). Significance values: ns: p > 0.05; *: p <=0.05; **: p <= 0.01; ***: p <=0.001; ****: p <= 0.0001. FAFB data built from FlyWire reconstructions.

We used early matching results for a total of 1,543 right hemisphere singletons to their homologs on the left hemisphere in the FlyWire data, which will be reported in an upcoming publication (Schlegel et al. 2023, in prep, see Methods). Because the HemiBrain data set is largely missing its left hemisphere, as well as large portions of the right one, we matched a smaller number (1,257) of right hemisphere FlyWire singletons to right hemisphere HemiBrain singletons. See Methods for how we calculate neuron-level transmitter predictions.

We found good agreement between left-right matched FlyWire singletons (Fig. 4b, left) and FAFB-HemiBrain matched singletons (Fig. 4c, left) for cholinergic, glutamatergic and GABAergic pairs. Inconsistent results between matched neurons were more common with singletons slated to express dopamine, serotonin or octopamine. The average confidence score (see Section 4.2.1) for FlyWire right (mean, 0.79, s.d., 0.16), FlyWire left (mean, 0.79, s.d., 0.17) and HemiBrain (mean, 0.67, s.d., 0.14) neurons was significantly higher (Fig. 4D) when there was no mismatch between our paired neurons, than when there was a conflict (FlyWire right: mean, 0.62, s.d., 0.22, FlyWire left: mean, 0.47, s.d., 0.14, hemibrain: mean, 0.43, s.d., 0.13). These scores represent how confident our network is in its prediction across all the presynapses of a given neuron (see Methods). Given the scores we see for well-matched neurons, we suggest that users of our resource to implement a stringent neuron-level transmitter prediction confidence score threshold of ∼0.62 and ∼0.53 for FlyWire and HemiBrain neurons respectively (one s.d. lower than the mean) if they need to be confident in the result for their neurons of interest.

We were curious as to whether matched singleton pairs correlate in their neuron-level transmitter prediction scores (Fig. 11a). When the network is less confident, it is consistently less confident for both members of a pair. This might indicate that the network has encountered a biological situation outside of our training data. For example, a different transmitter or co-expression situation. However, if it is less confident for one member of the pair, and much more confident in the other, this might be more indicative of differences in data quality. The similarity between matched pairs was greater among the FAFB-hemisphere matches than the FAFB-HemiBrain matches (Fig. 4e). This is likely because of inter-data set and model differences, since we can see strong correlations between our neuron-level transmitter prediction scores for our FlyWire left-right matched pairs (Fig. 4b right). Less confident singleton cell types may therefore differ in their biology from neurons in our ground truth data.

Similarly, we examined neurons grouped into isomorphic cell types in the HemiBrain data set (Bates et al., 2019; Scheffer et al., 2020a). We found that 8.0% of cell types (95 neuronal cell types) with more than one member per hemisphere did not have the same neuron-level transmitter prediction for each member. In these conflicted types, the neuron-level transmitter prediction score was significantly lower. Even more significantly, we matched 469 neuronal cell types between FlyWire and the HemiBrain data sets and found that > 95% agree in their modal neuron-level transmitter prediction between the two data sets (Fig. 11b). Again, the average prediction confidence score strongly correlated between the two data sets. Together, this suggests that there may be a biological factor, *e*.*g*., the expression of transmitters not in our training data, that leads to lower prediction scores and incorrect or inconsistent predictions with certain neuronal types, rather than a confound related to the EM data sets themselves.

### 2.7 Overview of transmitter expression in the *D. melanogaster* central brain

Having assessed the accuracy and consistency of our transmitter predictions, we next wanted to get an overview of transmitter usage in the *D. melanogaster* central brain. The central brain excludes the animal’s optic lobes, which are not present in the HemiBrain data set and are in the process of being reconstructed in FlyWire at time of writing. We examined all 11,277 whole neurons in the HemiBrain data set (see supplemental data file 7.1.3)., and 27,706 intrinsic brain neurons neurons that had been well reconstructed in FlyWire by the end of 2022, which we could “skeletonize” and “split” into a separable axons and dendrites (Schneider-Mizell et al., 2016) (see supplemental data file 7.1.4). We excluded Kenyon cells due to their complicated expression profile (see below). Our neuron-level transmitter predictions across our two neuron pools are relatively similar (Fig. 5a). Almost half the neurons in the HemiBrain are predicted to be cholinergic (FlyWire: 55.0%, HemiBrain: 45.7%) with similar fractions for glutamatergic and GABAergic neurons in both data sets (FlyWire: 24.3% glutamate versus 13.9% GABA, HemiBrain: 25.6% glutamate versus 17.0% GABA). Single cell RNA sequencing has suggested a similar break down across the central brain (44%:44% cholinergic, 23%:14% glutamatergic, 11%:15% GABAergic neurons, Davie et al. (2018) versus Croset et al. (2018)).

In *D. melanogaster*, acetylcholine acts as the principle excitatory transmitter and GABA the principle inhibitory transmitter. Glutamate can act in an either inhibitory or excitatory capacity but has only been reported as inhibitory in the central nervous system to date (Liu and Wilson, 2013; Lu et al., 2022; McCarthy et al., 2011; Molina-Obando et al., 2019). If we assume glutamatergic synapses to be inhibitory, then the ratio of inhibitory:excitatory neurons (FlyWire: 38.2%:55.0%, HemiBrain: 42.6%:45.7%) is close to parity in the HemiBrain. The largest discrepancy is in the proportion of predicted dopamine neurons (FlyWire: 18.5%, HemiBrain: 6.4%). This could be due to erroneous over-prediction of dopamine in the FAFB data set, though we may also simply have captured more dopaminergic neurons in our FlyWire neuron pool. Interestingly, if we look at how many automatically detected mitochondria (Plaza et al., 2022) exist per neuron, we can see that GABAergic neurons have slightly more mitochondria per micron cubed than cholinergic or glutamatergic neurons. Dopaminergic neurons have the most. Mitochondria ‘power’ cells, including neurons and their high abundance could indicate the level of energy use. Dopaminergic neurons have, though, the smallest somata (Fig. 14b), while serotonergic and octopaminergic neurons have larger nuclei and cell bodies.

In both data sets, we can see the proportions of neurons predicted with different confidence levels (Fig. 5b). Our neuron-level transmitter prediction confidences for the neuromodulators dopamine, serotonin and octopamine are not as good as those for the fast-acting transmitters. They make up a smaller proportion of the total neuronal inventory (FlyWire: 20.3%, HemiBrain: 8.2%) than the fast-acting transmitters. Because of this, the data set discrepancy with dopamine and because our matched singleton analysis (Fig. 4b and c) demonstrated that the network is more often mislead in its prediction for neuromodulators, we exclude them from some of the subsequent biological analyses, instead focusing on the fast-acting transmitters acetylcholine, GABA and glutamate. However, our prediction provide a starting place for looking for neurons with unusual transmitters or discovering, for example, unknown dopaminergic and serotonergic neurons, or assessing the pool of putative octopamine neurons from the literature (Busch et al., 2009) (Fig. 12d).

If we decompose the brain into its canonical brain regions for the HemiBrain data set where we have a dense reconstruction, we can see that the proportion of synapses of a given transmitter type varies between brain regions (Fig. 5d). For example, there is a lot more glutamate use in the fan shaped body, a region involved in navigation, whereas the primary source of inhibition for the ellipsoid body, a linked region involved in navigation, is GABA. Since dopamine has been heavily implicated in many forms of synaptic plasticity, its high proportion in certain neuropils may indicate that these regions experience more synaptic plasticity than others, and are therefore sites of learning within the connectome.

We note that our network’s tendency to incorrectly predict Kenyon cells as dopaminergic means that we should treat this analysis with caution. RNA sequencing data has shown that Kenyon cells express the genes necessary for cholinergic transmission (Barnstedt et al., 2016; Crocker et al., 2016; Croset et al., 2018). Despite this, our neuron-level transmitter predictions guess dopamine across all Kenyon cell types (FlyWire: 99.9%, HemiBrain: 99.9%) with high confidence (FlyWire: mean, 0.627, s.d., 0.99, HemiBrain: mean, 0.54, s.d., 0.046). The *αβ* and *γ* Kenyon cells are known to express the short neuropeptide F precursor, which may indicate neuropeptide transmission. In addition, *Ddc*, which encodes a protein responsible for converting L-DOPA to dopamine, is expressed in *α*^′^ *β*^′^ and *γ* Kenyon cells, where it could play a role in the biosynthesis of dopamine or another aromatic L-amino acid (Croset et al., 2018). Co-expression with transmitters outside of our training data may be one major reason why our results can sometimes differ from what we expect, given wet-lab investigations. A further reason may be due to contamination by nearby presynapses belonging to other, truly dopaminergic neurons. The cube of image data (edge length 640 nm) centered on each predicted presynaptic location was not masked with neuronal identity, and it is possible that features from proximal presynapses have skewed the result (see Methods for further discussion). Many mushroom body dopaminergic PPL1 and PAM neurons were used in our ground truth data, and indeed most presynapses in the highly synapse-dense region of the mushroom body are predicted as dopaminergic (Fig. 5d).

Co-expression of transmitters, especially fast-acting transmitters alongside neuropeptides and other neuromodulators, is expected to be common in the brain although examples at the resolution of individual cell types is sparse. A singleton cell type, the dorsal paired medial, DPM, neurons, have been found to express the fast-acting transmitter GABA, the neuropeptide amnesiac and the monoamine serotonin. They were not used in our ground truth data, and in both data sets are predicted to be dopaminergic. The neurons typed are thought to express both dopamine and serotonin (Mao and Davis, 2009), but we predict them only as dopaminergic. The hDelta cell types are a set of 190 neurons whose morphologies are very similar, and who segment the fan-shaped body of the fly. They have been popular for investigation in recent years (Lu et al., 2022). However, our HemiBrain prediction results estimate 33% to be cholinergic and 42% to be dopaminergic. While the field lacks authoritative data, we would have expected them all to express the same transmitter or set of transmitters, most likely acetylcholine. In cases like this, it is difficult to assess from our results alone whether these cell types express acetylcholine and/or dopamine, and/or other transmitters as was the case with DPM. Fortunately it is not common, only 8.0% of cell types in the HemiBrain have a transmitter prediction conflict.

When the network experiences something outside of its learning set, it is usually less confident in its erroneous prediction. For HemiBrain neurons in our ground truth, the mean neuron-level transmitter prediction confidence was 0.57, s.d., 0.17. For neurons we know to use transmitters outside of the six we learned, we found that glycinergic neurons (Frenkel et al., 2017) were mostly predicted to use acetylcholine (mean confidence 0.66), Allatostatina (Frenkel et al., 2017) were mostly predicted to use GABA (0.7) (even though it co-transmits with glutamate (Croset et al., 2018)), Corazonin (Kubrak et al., 2016) use was predicted as acetylcholine (0.3), Drosulfakinin (Scheffer et al., 2020a) use was mostly predicted as octopamine (0.47), insulin (Scheffer et al., 2020a) use was mostly predicted as acetylcholine or serotonin (0.44), IPNa (Shafer et al., 2006) use was mostly predicted as glutamate (0.32), NPF (Scheffer et al., 2020a) use was mostly predicted as octopamine (0.46) and SIFamide (Scheffer et al., 2020a) use was mostly predicted as octopamine (0.43). Across the board, for HemiBrain neurons in our ground truth, the mean neuron-level transmitter prediction confidence was 0.57, s.d., 0.17.

### 2.8 Distribution of Neurotransmitter Predictions within Developmental Units

The nervous system may already effectively “group” neurons by their transmitter expression (Lacin et al., 2019). The neurons of the central nervous system are generated by a set of stem cells known as neuroblasts. During division neuroblasts generate two cells, one additional stem cell and one cell that further divides into two sibling neurons. In only one of these siblings the so-called Notch pathway is activated, leading to two different *hemilineages* of neurons within each lineage (Kumar et al., 2009; Truman et al., 2010; Sen, 2019; Lacin et al., 2019) (or more than two in the case of Type II neuroblasts, which mainly contribute to the central complex) (Fig. 6a).

**Figure 6:**
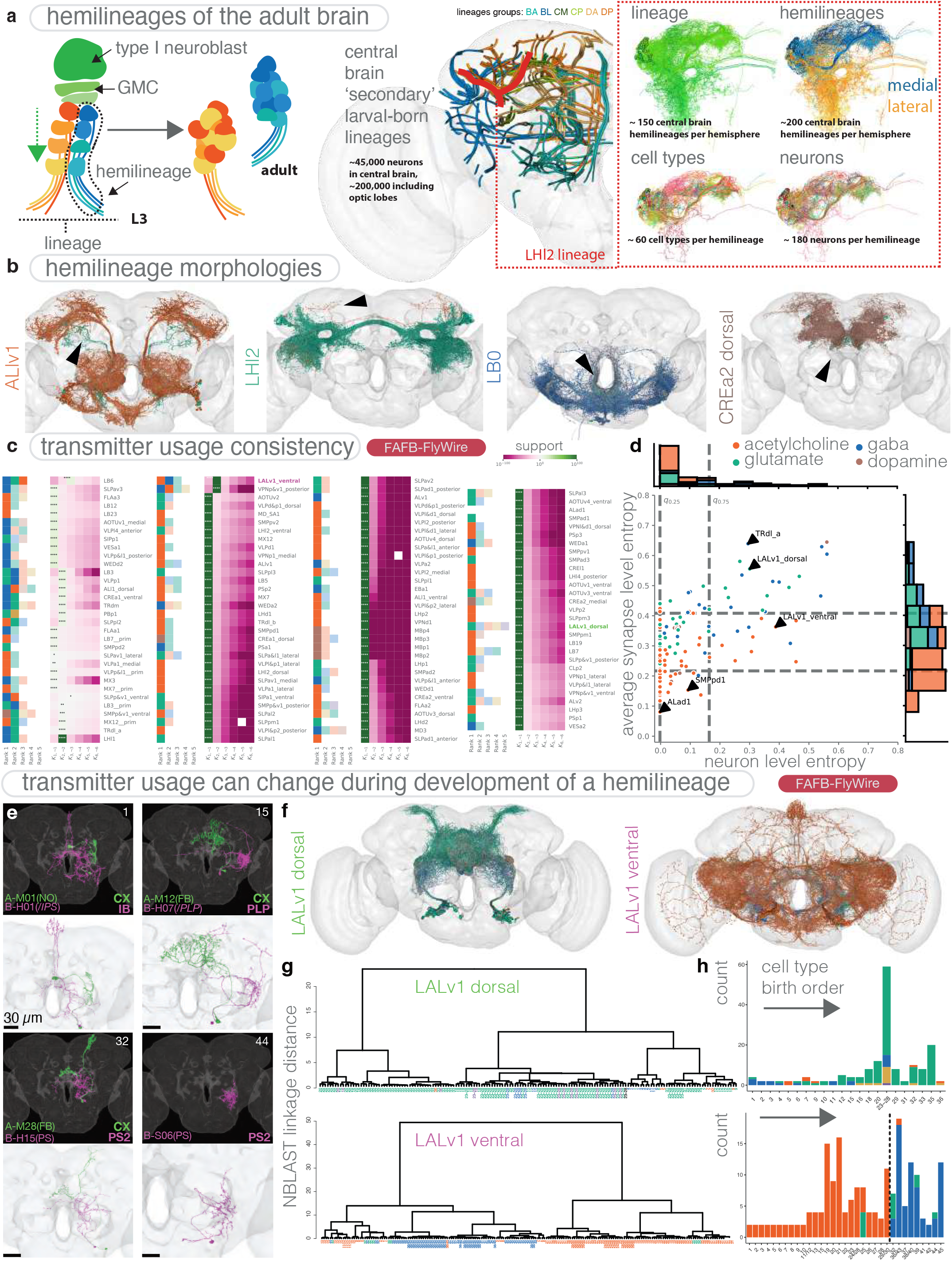
**(a)** Illustration of the development of Type I hemilineages in the adult *D. melanogaster* brain. Illustration shows (left) the progression of a Type I neuroblast from third-instar (L3) larva into the adult, GMC, ganglion mother cell and (right) breakdown of a single secondary lineage, LHl2 (also known as DPLal2) into its two constituent hemilineages. Neuronal reconstruction data from FAFB-FlyWire shown, demonstrating how a lineage of neurons can be distilled into smaller and smaller groups: hemilineages, cell types and individual neurons. Each lineage, hemilineage and cell type exist in duplicate, one copy on each side of the nervous system, effectively providing an internal control for validating our predictions. **(b)** Example images of neurons from the FAFB-FlyWire in four different hemilineages, meshes coloured by neuron-level transmitter prediction. A black arrow in each image points to one stray member of the hemilineage, with a different neuron-level transmitter prediction and gross morphology. These could be first-born neurons from the hemilineage, which often have aberrant morphology (Lee et al., 2020). **(c)** Bayes factor analysis of hemilineage consistency in the adult *D. melanogaster* brain. For each hemilineage row, the right set of columns corresponds to the likelihood ratio of the hemilineage expressing that number of transmitters versus the likelihood of any other number 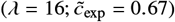. Evidence strength: *: *K* ≥ 10^1/2^; **: *K* ≥ 10^1^; ***: *K* ≥ 10^3/2^; ****: *K* ≥ 10^2^. (Jeffreys, 1998). The left set of columns indicate the frequency ranked transmitter predictions within the hemilineage, with ranks greater than the maximum likely number of transmitters shaded lighter. Data ordered by clustering rows by their Bayes factors. **(d)** Neuron level entropy (*H N*_*h*_) vs average synapse level entropy (*H* (*S*_*h*_)) for all predicted hemilineages with more than 10 neurons and more than 30 synapses per neuron (**a**). Dashed lines indicate 25% and 75% percentiles: *q*_25_ (*H* (*N*_*h*_)) = 0.00, *q*_25_ (*H* (*S*_*h*_)) = 0.17 and *q*_75_ (*H* (*N*_*h*_))= 0.22, *q*_75_ (*H* (*S*_*h*_))= 0.41. **(e)** Examples of neurons from the FAFB-FlyWire data set (lower panels, white background) matched to light-microscopy single neuron images (higher panels, dark background) from Lee et al., (2020). Neurons in green are from the Notch-ON hemilineage of LALv1 (LALv1 dorsal); and neurons in magenta are from the Notch-OFF hemilineage (LALv1 ventral). They were matched with a combination of visual inspection and (C) morphological clustering with the NBLAST algorithm (Costa et al., 2016). **(f)** Images of neurons from the FAFB-FlyWire data set from the lineage LALv1. This lineage has two hemilineages (see example in A), dorsal and ventral. Our Bayes factor analysis highlights LALv1 ventral as a multi-expression hemilineage (see coloured labels in C). Neuron meshes coloured by neuron-level transmitter prediction. LALv1 is also, fortunately, one of few hemilineages whose birth order has been delineated (Lee et al., 2020). **(g)** Dendrograms generated based on morphological clustering of individual neurons, and the nodes are coloured by the predicted neurotransmitter of that neuron. The node labels represent the matched birth order in Lee et al., 2020 (larger number = later born). Notably, in LALv1 ventral, neurons with different morphologies (i.e. in different dendrogram clusters) have relatively well-separated transmitter profiles. **(h)** Histograms of neuron-level transmitter prediction by birth order and hemilineage. Notably, there is a switch from acetylcholine to GABA from group 29/30 to group 32 for the LALv1 ventral hemilineage, indicating that there may be a switch of transmitter expression during development. FAFB data built from FlyWire reconstructions.

Neurons can be born in this way during embryogenesis are known as primary neurons (∼10% of the adult brain (Marin et al., 2005; Veverytsa and Allan, 2013) but may also account for a majority of neuromodulatory neurons (Dacks et al., 2006; Busch et al., 2009)) and those made during larval development as known as secondary neurons. Only secondary hemilineages (each ∼∼90 neurons) can easily be demarcated in adult *D. melanogaster*. Neuronal fibers co-fasciculate with neurons from the same hemilineage and form bundles as they enter the neuropil from the insect brain’s outer-layer of cell bodies. These have been mapped by light microscopically and assigned to discrete lineage clones (Ito et al., 2013; Yu et al., 2013; Wong et al., 2013; Lovick et al., 2013). These *hemilineage-associated tracts* can also be readily identified in EM stacks based on their point of entry and subsequent trajectory. It should be noted that cases have been identified (Lee et al., 2020) where a single hemilineage-associated tract is composed of two hemilineages. Since such cases are rare, we will in the following consider hemilineage-associated tracts as fiber units belonging to one hemilineage.

Lacin et al. (2019) showed that each hemilineage in the ventral nervous system uses just one of the fast-acting transmitters, acetylcholine, glutamate or GABA, even though mRNA transcripts for combinations of these can appear in the nucleus (Lacin et al., 2019). We refer to this principle as *Lacin’s law*. This raises the question whether the same holds true in the adult brain.

Using the presented classifier, we predict the neuron-level transmitter prediction of all neurons in 161 selected brain hemilineages, out of a total of ∼231 per hemisphere. These hemilineages were chosen on the basis on our annotation status in the FlyWire data set at the time of writing (shared for early access by Schlegel et al., in prep). We only included hemilineages for which we could find a similar number of neurons on the left and right hemispheres after multiple rounds of review, and whose gross morphology matched extant light-level clonal data (Ito et al., 2013; Yu et al., 2013; Wong et al., 2013; Lovick et al., 2013) (see supplemental data file 7.1.5). Our mean difference in neuron number between the left and right hemisphere copies was only 3. How mixed a hemilineage is in its neuron-level transmitter predictions also correlates strongly been to the left and right hemisphere copies (Fig. 13c). The majority of our predictions show homogeneity of neurotransmitter identity within a single hemilineage, in line with Lacin’s law (Lacin et al., 2019) (Fig. 6b,c, Fig. 13a). Two hemilineages, CREa1 dorsal and CREa2 dorsal, are known to produce majority dopaminergic neurons (Lee et al., 2020), the PAM dopaminergic neurons that innervate the mushroom body and play a role in associative learning and memory (Aso et al., 2014) and dopaminergic neurons of the fan-shaped body (Hulse et al., 2021). We see this clearly in our neuron-level transmitter predictions for these hemilineages (Fig. 13a).

However, some hemilineages present mixed expression (Fig. 13a,b). In some cases a small number of neurons deviate from the majority prediction (Fig. 6b). These neurons may be the “first-born” neurons of the hemilineage, which have been shown to often have a deviant morphology to the rest of the hemilineage (Lee et al., 2020). Our results suggest that they may also be deviant in their gene expression. In some other hemilineages larger groups of neurons deviate (Fig. 6c, Fig. 13a), and in these cases neuron-level transmitter predictions seem to break down by morphological subgroup within the hemilineage (Fig. 13D). For example, DL1 dorsal contains both glutamatergic neurons that project to the fan-shaped body, and cholinergic ones that do not (see supplementary image data). LALv1 ventral contains cholinergic neurons with lateral arbours in the ventral brain, and GABAergic neurons that are placed medially (Fig. 6e,f). In other cases, such as that of LHl1, ALv2, LHl2 and VLPl4 dorsal, hemilineages can show mixed expression despite gross morphological similarity within each hemilineage (Fig. 13a). To explore why, we mapped the two hemilineages in LALv1 to a recent analysis that delineated the birth order of LALv1 neurons in time (Lee et al., 2020). We found that for LALv1 ventral, there is a switch in transmitter expression after the 31st neuron is born (Fig. 13B,D), and that these neurons look morphologically different (Fig. 13c). The birth order for others of our mixed hemilineages is unknown. However, we suggest that is is likely that some hemilineages experience a shift in transmitter usage due to a switch in transcription factor expression in later born neurons. This may also incur a correlated gross morphological change.

We next asked how likely it is to observe a given prediction of transmitters in a hemilineage under some error rate given by the confusion matrix on the test set, and the assumption that all neurons in the hemilineage have the same underlying transmitter. We can then compare this likelihood to the alternative hypothesis that a hemilineage consists of neurons with more than one transmitter.

We calculate the Bayes factor 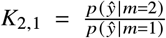 and 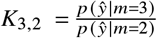 for our selected hemilineages, that is, the likelihood ratio of the model’s predictions given a hemilineage expresses 2 rather than 1 transmitters or 3 rather than 2 transmitters, respectively (Fig. 13d, Fig. 13e). For LALv1 ventral and 15 other hemilineages there is decisive evidence (*K*_2,1_ ≥ 10^2^) for the presence of two distinct transmitters rather than one for a large range of expected accuracies 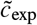 (Fig. 13e). However, some of these hemilineages such as TRdla show high synaptic entropy *H* (*S*_*h*_) (Fig. 6D), indicating that individual neurons within the hemilineage contain substantial multimodal transmitter predictions. As such, multimodality at the neuron level is at least partially explained by uncertain or inhomogeneous predictions between individual synapses within a neuron. This is in contrast to hemilineage LALv1 ventral, which has a synaptic entropy within the 75% percentile. In these hemilineages, large Bayes factor *K*_2,1_ values directly stem from neuron level segregation of the predicted transmitters within the hemilineage.

Maximal one-versus-rest Bayes factors (*K*_*m*,¬*m*_) summarize our predictions of the number and set of transmitters for each hemilineage (Fig. 6C). We found 12 hemilineages with decisive evidence (*K* ≥ 10^2^) of expressing two transmitters, one with very strong evidence (*K* ≥ 10^3/2^), and one with substantial evidence (*K* ≥ 10^1/2^). One hemilineage showed decisive evidence for all three fast-acting transmitters, while one showed substation evidence (*K* ≥ 10^1/2^). We predict the remaining 109 hemilineages to express a single transmitter (n=107 decisive, n=1 strong, n=1 substantial).

### 2.9 Distribution of Neurotransmitter Usage Between Axons and Dendrites

In addition to having predictions for each neuron’s synapses and its transmitter type, we can also use the neuron’s morphology to estimate which of its synapses belong to its axon and which to its dendrite (Fig. 7a) (Cajal, 1911; Rolls, 2011). For each neuron in our pool (Fig. 5a), we computed the axon and dendritic compartment using a graph theoretic algorithm developed in Schneider-Mizell et al. (2016). We could then ask whether presynapses of different transmitter types differ in their distribution across axons and dendrites, and whether they are enriched or not at certain connection types. It is important to note that only a neuron’s presynapses (its outputs) determine its transmitter usage. Postsynapses (its inputs) receive transmitter released at the presynapses of upstream neurons. The types of transmitter received can be very diverse.

**Figure 7:**
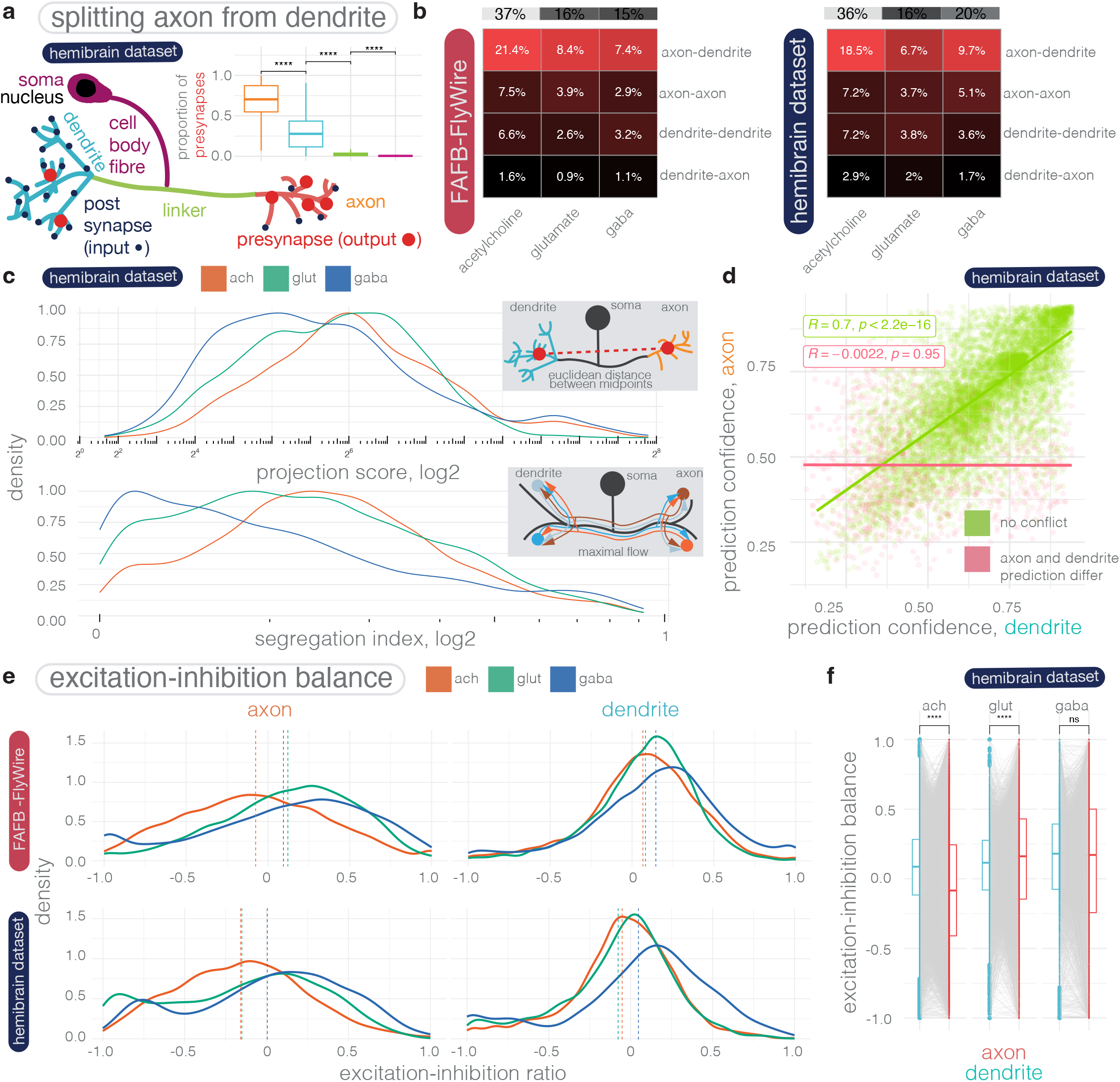
Comparing neurons’ transmitter usage predictions between connectome data sets (FAFB-FlyWire and HemiBrain) and between hemisphere in the same data set (FAFB-FlyWire). **(a)** Schematic of a neuron broken into its separate neuronal compartments. In general, the soma, cell body fibre tract and linker cable (primary dendrite) do not contain large numbers of synaptic contacts. Inset, the proportion of presynapses in each of the four compartment types. The raw numbers of synapses after filtering can be quite different between the FAFB-FlyWire and HemiBrain data sets (FAFB-FlyWire: median 202, hemibrain: median 386). **(b)** How the connectome’s synaptic budget is spent, across different connection types. Heatmaps for the FAFB-FlyWire (left) and HemiBrain (right) data sets show the proportion of synaptic contacts from neurons of different predicted transmitter types (columns) used in different inter-compartmental connection types (rows). Only neurons with a distinguishable axon and dendrite used in this analysis, FAFB-FlyWire: 9,123, hemibrain: 10,122 neurons. **(c)** Scaled density plots showing neuronal polarity by neuron-level transmitter prediction. Upper, distributions of projection score across neuron pool. The projection score is the distance in Euclidean space between the midpoint of the predicted dendrite and the axon. Segregation index is a metric for how polarised a neuron is; the higher the score the more polarised the neuron (Schneider-Mizell et al., 2016). FAFB-FlyWire: 9,123, hemibrain: 10,122 neurons. **(d)** Correlation between compartment-level transmitter prediction score for axons and dendrites. Each point is a separate neuron in the HemiBrain data set, N = 10,122, 11.0% disagree on the compartment-level transmitter prediction (red). Scatter plot displays the Pearson’s product-moment correlation, giving R, the coefficient and associated p-value. **(e)** Scaled density plots showing the distribution of excitation-inhibition (EI) balance (proportion of excitatory input minus proportion of inhibitory input) across neuron-level transmitter predictions and compartments. Neurons have been pooled into three groups by their transmitter identity, colours. Upper, FAFB-FlyWire, lower, HemiBrain data set. Dotted lines indicate the mean value calculated across all neurons. The more positive the balance, the more excitation a neuron receives. Note neurons are unmatched between data sets in this analysis, *i*.*e*., the corpus of neuronal cell types from each data set is partially overlapping but distinct, cell type content my account for some of the FAFB–HemiBrain differences. **(f)** Comparison of EI balance between compartments in the HemiBrain data set. Data compared using Wilcoxon two-sample tests. Significance values: ns: p > 0.05; *: p <=0.05; **: p <= 0.01; ***: p <=0.001; ****: p <= 0.0001. FAFB data built from FlyWire reconstructions.

A majority of presynapses are found on the axon (flyWire: median 76%, s.d., 21%, hemibrain: median 70%, s.d., 21%). However, *D. melanogaster* neurons often have a large proportion of their output sites on their dendrites (flyWire: median 23%, s.d., 21%, hemibrain: median 30%, s.d., 21%). Few presynapses are to be found on the cable leading from the cell body, or connecting axon and dendrite (Fig. 7a). The proportion of presynapses spent on the axon versus the dendrite differ slightly between neurons of different neuron-level transmitter predictions (Fig. 14h).

Between neurons, axons and dendrite can connect to one another with either being the source or the target (Fig. 7b). In the brain, the majority of the synaptic budget is spent on axodendritic connections (flyWire: 55%, hemibrain: 48%). However, a large fraction is spent on axo-axonic connections (flyWire: 22%, HemiBrain: 20%) and similar amount on dendro-dendritic connections (flyWire: 19%, hemibrain: 21%). These figures are comparable to those recently reported for the *D. melanogaster* larva (axo-dendritic: 54%, axo-axonic: 36%, dendro-dendritic: 8%, dendro-axonic: 3%), which generated by the same approach to separate axon from dendrite (Winding et al., 2022; Schneider-Mizell et al., 2016). Cholinergic input to dendrites is three-fold larger than to axons, while the difference for glutamatergic or GABAergic connections is about two-fold Fig. 7b), suggesting that a key role for GABAergic and glutamatergic neurons is to provide inhibition to axons.

We were interested to ask whether the degree of axon-dendrite polarization differs between neurons with different neuron-level transmitter predictions (Fig. 7c). GABAergic neurons axons and dendrites are closer together in Euclidean space (Fig. 7c upper). In addition, they have a lower segregation index (Schneider-Mizell et al., 2016) than cholinergic or glutamatergic neurons (Fig. 7c lower), and are smaller in terms of cable length (Fig. 14a). This shows that on average there are gross morphological differences between even the two putative inhibitory classes, GABAergic and glutamatergic neurons. The difference in segregation index is largely due to a greater number of presynapses per unit of cable in GABAergic neurons Fig. 14c). The number of inputs or outputs per unit cable length in other compartments is more similar.

We were also curious as to whether axons and dendrites could receive different compartment-level transmitter predictions in their axon compared with their dendrite. In the FlyWire data set, but not in the HemiBrain, synapse-level transmitter prediction scores were higher for axons than for dendrites (Fig. 14d). Typically, we do not expect these compartments to use different transmitters. We found a small percentage of axon-dendrite prediction mismatches (flyWire: 6.5%, hemibrain: 11.0%) (Fig. 7d, Fig. 14i). A majority of these mismatches were dopamine or glutamate predictions for the axon, and acetylcholine for the dendrite (Fig. 14i). In neurons where there was no mismatch, there was a strong correlation between the compartment-level transmitter prediction for the axon and the dendrite (Fig. 7d), suggesting that the local image features impacting our prediction could be the same for both (even though axon and dendrite might be far from one another in Euclidean space, Fig. 7c upper). For mismatches, there was no correlation. In remains to be seen in wet-lab experiments whether some neurons could use different transmitters between compartments.

Assuming that glutamate has an inhibitory action, we can calculate the excitation-inhibition (EI) balance across our two neuron pools (Fig. 7e, Fig. 14d). The distribution for dendrites of acetylcholine glutamate and GABA neurons looks normal and is centred just positive of 0 (median, -0.02) but the distribution for EI balances for axons differ. For cholinergic neurons it is a wider normal distribution centered slightly more negatively (median, - 0.16). For GABAergic and glutamatergic neurons there are two peaks, a negative and a positive peaks. In particular, the EI balance for GABAergic axons was skewed negative (Fig. 7f). This indicates that some neurons receive a lot of inhibition to their axons, while others receive a lot of excitation possibly as a part of feed-forward inhibition or excitation motifs. There is a weak correlation between the EI balance for the axon and the dendrite, which is a little stronger in GABAergic and glutamatergic neurons (Fig. 7e). Breaking neurons down further by Strahler order (Fig. 14f) we did not see an obvious difference in the distribution of inputs of different neuron-level transmitter prediction Ė I balance did not correlate with total neuron size, dendrite size, axon size, presynapse counts or postsynapse counts (data not shown).

### 2.10 Neurotransmitter Identity of Synaptic Inputs

Each neuron can receive a diverse range of inputs. The majority of inputs are weak, below five connections between the source (upstream) and target (downstream) neuron, with GABAergic connections proving slightly stronger on average (Fig. 8A). The distribution of input by transmitter type along the neuronal arbour is not obviously skewed across our whole pool of neurons (Fig. 8B), though particular neurons have been reported to exhibit physiologically relevant strategic synapse placement (Felsenberg et al., 2018b).

**Figure 8:**
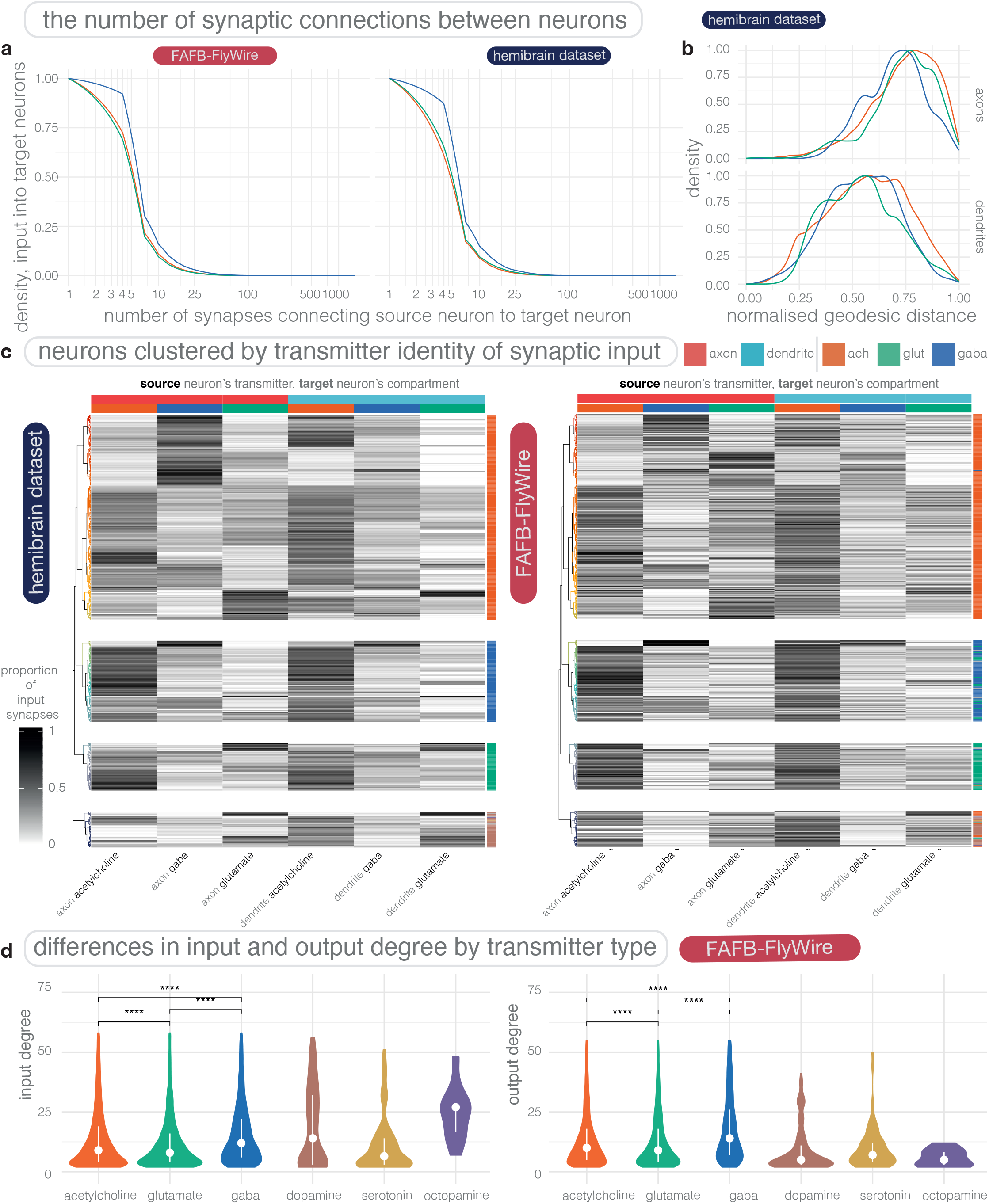
Input profiles for brain neurons based on inputs’ transmitter identity and location. **(a)** Scaled density plot showing the density of input connections onto all FAFB-FlyWire and HemiBrain neurons (facets) at different synaptic weights (X-axis, log2). **(b)** Scaled density plots showing the max-normalised geodesic distance (the distance along a neuron’s arbour) from input synapses (coloured by the source neuron’s neuron-level transmitter prediction) to the target neurons’ cell body. **(c)** Equivalent 451 neuronal cell types from the FAFB-FlyWire and HemiBrain data sets clustered by input type. Only neurons for which at least 50%of inputs came from well reconstructed and predicted neurons in our 27,706 FAFB-FlyWire neurons or 11,277 HemiBrain neurons were used. For each source neuron to target neuron connection, we used the identity (neuron-level transmitter prediction), location (neuronal compartment) and normalised connection weight (number of synaptic contact made on that compartment / total number of synaptic inputs to the target neuron). We calculated cell type averages, and separated target cell types by their transmitter prediction and then clustered within each grouping. Heatmaps show the proportion of synaptic input onto the axon (upper horizontal color bar, red) and dendrite (blue), separated by the neuron-level transmitter prediction for each input (lower horizontal color bar). Each row is a separate neuronal cell type (see Fig. 15C for names). Cell types are grouped by a hierarchical clustering within their neuron-level transmitter prediction class (vertical color bar: acetylcholine, glutamate or GABA) employing Ward’s clustering criterion. This clustering was performed in the HemiBrain data set and applied to the FAFB-FlyWire data set. Dendrogram (left) colors show a split into 30 groups. The same dendrogam is used in both heatmaps. Cosine similarity, z = 0.892, p-value ¡ 0.0001, 100,000 row shuffles. **(d)** Differences in the number of outgoing and incoming connections by neuron-level transmitter prediction. The input and output degree for a neuron is the number of unitary connections it has incoming and outgoing, respectively (the number of synaptic pairs, regardless of synaptic weight). All source-target connections with a synaptic count ≥ 10 included. Left, box plots show the distribution of input degrees by the target neurons’ neuron-level transmitter prediction. Right, output degrees by the source neurons’ neuron-level transmitter prediction. Violin plots show the median (dot), the first and third quartiles (lines, the 25^th^ and 75^th^ percentiles). The data out of the bounds of the 5^th^ and 95^th^ percentiles have been removed for ease of visualization. Significance values: ns: p > 0.05; *: p <=0.05; **: p <= 0.01; ***: p <=0.001; ****: p <= 0.0001. FAFB data built from FlyWire reconstructions.

We can break down the input to a target neuron by the source neuron’s neuron-level transmitter prediction and compartment, *i*.*e*., axon versus dendrite. We wanted to know whether after doing this, the input profile of neuronal cell types that we have matched up between the HemiBrain and FlyWire (Fig. 11b) are the same, and how diverse they are.

We found that cell types cluster into distinct groups with different input profiles (Fig. 8c). Each cluster typically receives a large proportion of axo-dendritic cholinergic input, however the degree of other input types can vary widely. Groups tended to receive either strong acetylcholine, glutamate or GABA drive to their axons (Fig. 15a). The negative correlation between different types of transmitter input is weaker in the dendrite than for axons (Fig. 8b). The cosine similarity between the two matrices is high (> 0.9) indicating that the input profile of a cell type is similar between brains.

Interestingly in our FlyWire data, when grouping neurons by hemilineage, we can see that while each hemilineage can contain neurons from a range of clusters particular hemilineages can be enriched with neurons from certain clusters (Fig. 15b). This suggests that neurons that are born and develop together, expressing a similar gene profile receive similar types of inputs. Morphologies from a hemilineage can be diverse, and so this preference could be enforced by molecular targeting strategies as well as simply neurons’ physical location in space.

We were also interested to ask how many synaptic partners neurons of different neuron-level transmitter prediction have. We determined input and output degree for FlyWire (Fig. 8c) and HemiBrain neurons using a threshold of 10. In our HemiBrain data, cholinergic, glutamatergic and GABAergic neurons output to a similar number of target neurons (acetylcholine:, median, 9, IQR, 15, glutamate: median, 13, IQR, 21, GABA: median, 12, IQR, 28). Their input degrees are very similar to their output degree (acetylcholine:, median, 9, IQR, 17, glutamate: median, 12, IQR, 21, GABA: median, 12, IQR, 26). In our FlyWire data, GABAergic neurons more clearly had a greater input (acetyl-choline:, median, 9, IQR, 17, glutamate: median, 8, IQR, 13, GABA: median, 12, IQR, 19) and output degree (acetylcholine:, median, 10, IQR, 14, glutamate: median, 8, IQR, 14, 21, GABA: median, 16, IQR, 25) than cholinergic and glutamatergic neurons. Dopaminergic neurons target a smaller number of neurons, and receive input from a greater number of neurons (Fig. 8c right), in agreement with recent work exploring dopaminergic neurons of the mushroom body in Catmaid (Otto et al., 2020). For octopaminergic have the largest input degree, possibly because they tend to be larger (Fig. 14a).

### 2.11 Neurotransmitter use through layers of the olfactory system

Having examined transmitter usage from the perspective of individual neurons, we were interested to ask how transmitter use might differ across a neuronal circuit. Meta data for full networks of neurons have not yet been described for entire sensory systems within the FAFB data set. However, in recent work (Schlegel et al., 2021) we delineated the organisation of the *D. melanogaster* olfactory system as captured in the HemiBrain connectome (Fig. 9A). We distilled the HemiBrain olfactory system into is constituent cell types and classes. We had previously found that olfactory projection neurons receive a large amount of inhibition to their axons from local third-order inhibitory neurons (Bates et al., 2020b), a finding that we capture again here by examining the EI balance for different olfactory classes (Fig. 9B).

**Figure 9:**
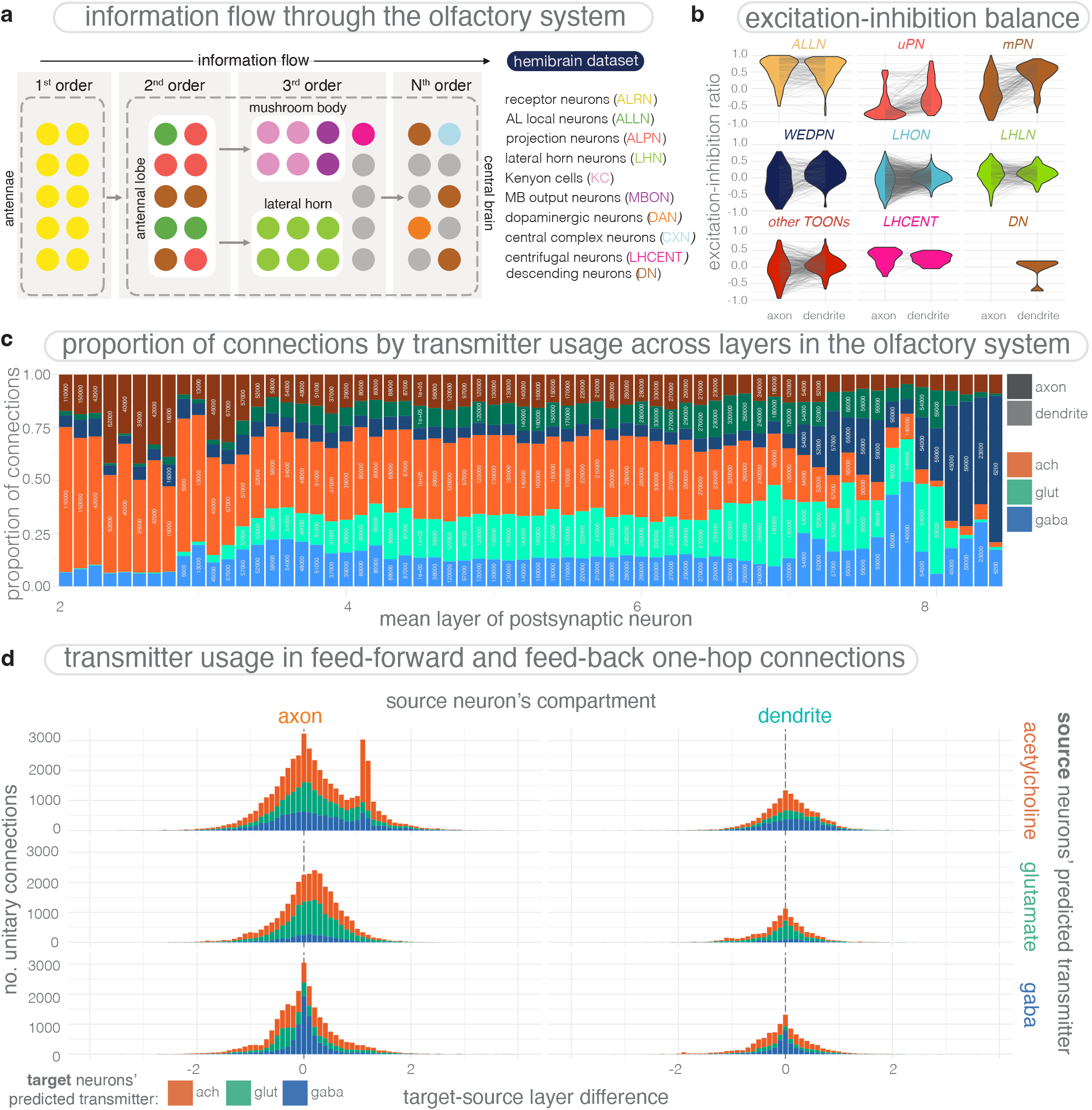
Transmitter use through layers of the olfactory system. **(a)** Schematic from Schlegel et al. (2021) depicting the organisation of the *D. melanogaster* olfactory system. **(b)** The excitation-inhibition (EI) balance across commonly known cell types of the olfactory system, described in Schlegel et al. (2021). Violin plots have superimposed lines connecting axons and dendrites that belong to the same neuron. **(c)** Transmitter input and compartment distribution through layers of the olfactory system. Plot shows a normalised stacked bar chart, binned by target neurons’ layer score, bin width 0.1. Bars colors indicate the source neurons’ neuron-level transmitter predictions and the hue indicates postsynapse location on target neurons, dark for axon, light for dendrite. Text shows the number of connections made onto target neurons in the given bin. Number of neurons: 24,549. Number of connections: ∼44,000,000. **(d)** Feed-forward and feed-back connectivity across the olfactory system by neuron-level transmitter prediction. Plot is faceted by source neuron-level transmitter prediction, colors indicate target neuron-level transmitter prediction. For each immediate unitary neuron connection between a source and target neuron we calculated a target-source layer difference, which is simply the layer value (Schlegel et al., 2021) for the target neuron - the layer value of the source neuron. The more positive the score, the more feed-forward the connection within the olfactory system. The more negative the score, the more feed-back the connection. The Y axis gives the number of unitary connections in each bin, bin width 0.1. A number of unitary connections is the number of synaptic pairs, regardless of synaptic weight, after a threshold 100 was applied. Histograms show the distribution.

In our prior work (Schlegel et al., 2021), we employed a simple probabilistic graph traversal model to “step” through the olfactory system, from sensory input neurons through to all good neurons in the HemiBrain data set (24,549), and record the position at which each neuron is encountered. We call the positions established by this procedure “layers”. These layers correspond to the mean path length from sensory neurons to any other HemiBrain neuron taking account of connection strengths over 10,000 runs.

Here, we show that as we progress through layers of the olfactory system, our input connection budget fluctuates (Fig. 9C). Initially, the majority is spent on cholinergic connections onto axons and dendrites. Progressing into the deeper layers, putative inhibitory connections become more common, both onto axons as well as dendrites.

Finally, we asked how neurons with different neuron-level transmitter predictions are represented in feed-back and feed-forward connectivity, through the olfactory layers we had described (Schlegel et al., 2021) (Fig. 9D). We found that outgoing connections from both a source neuron’s axon and dendrite were slightly biased towards targeting a neuron in a higher layer when that source neuron is cholinergic (mean 0.21, s.d., 0.75) or glutamatergic (mean 0.07, s.d., 0.57). However, GABAerigc neurons exhibit a more normal distribution (mean -0.02, s.d., 0.61). In aggregate, a larger proportion of cholinergic neurons (54%) and glutamatergic (56%) fed-forward through the olfactory layers, while GABAergic neurons were split 50:50, feedback:feed-forward.

### 2.12 Data Availability

We have made our results available via popular connectome data hosting services as well as direct downloads. Our transmitter classification network and associated training and prediction code (Synister) is available at https://github.com/funkelab/synister, as well as instructions how to access the FlyWire and HemiBrain predictions. Our synapse-level transmitter prediction and neuron-level transmitter predictions are hosted by extant connectome annotation and browsing services (HemiBrain: https://neuprint.janelia.org/?dataset=hemibrain:v1.2.1, flyWire: https://codex.FLywire.ai). The FlyWire synapse-level predictions can also be directly downloaded under https://www.dropbox.com/s/qv1rzsqekgj1zij/synister_fw_mat571_t11_synapses.feather?dl=0 and the HemiBrain predictions are available for download under https://storage.googleapis.com/hemibrain/v1.2/hemibrain-v1.

2-tbar-neurotransmitters.feather.bz2. We have provided the studies we have used to generate our ground-truth data (supplemental data file 7.1.1), identifiers for the neurons we used for our ground truth data (supplemental data file 7.1.2), our neuron-level transmitter predictions for each complete neuron in the HemiBrain data set (supplemental data file 7.1.3) and FlyWire data set (supplemental data file 7.1.4) and images of each of our chosen brain hemilineages with neurons coloured by their predicted transmitter expression (supplemental data file 7.1.5) as supplemental data for this paper.

## 3 Discussion

A consistent criticism of large scale connectomic efforts is that they struggle to make verifiable, functional predictions for circuit mechanisms. Machine learning tools already underpin recent progress in the field, *e*.*g*., to segment neurons (Berning et al., 2015; Funke et al., 2018; Januszewski et al., 2018; Lee et al., 2019; Sheridan et al., 2022; Schmidt et al., 2022; Dorkenwald et al., 2022; Svara et al., 2022) or to detect synapses (Buhmann et al., 2018; Kreshuk et al., 2015; Staffler et al., 2017; Buhmann et al., 2019; Ryan et al., 2016; Dorkenwald et al., 2017; Svara et al., 2022). Here we showed that machine learning methods can not just be used to automate tedious reconstruction tasks, but can also predict functional properties from ultrastructure alone: By amalgamating our new synapse-level transmitter predictions into neuron-level transmitter predictions, it is possible for us to generate a comprehensive transmitter wiring diagram for the connectome of *D. melanogaster* that includes a prediction of the *sign* of each connection, *i*.*e*., whether the upstream neuron excites, inhibits, or otherwise modulates a downstream target. This circumvents a major bottleneck in transmitter identification in neurobiological circuits, *i*.*e*., matching light microscopy expression patterns to connectomic neuronal reconstructions (Bates et al., 2019).

We have made our results in the brain (the FlyWire and HemiBrain data sets) publicly available (see Section 2.12). They have already been used by several studies (Lu et al., 2022; Baker et al., 2022; Schlegel et al., 2021; Li et al., 2020a; Engert et al., 2022; Westeinde et al., 2022; Eichler et al., 2023) in order to generate neurobiological hypotheses. Our results are therefore a significant step forward in increasing the interpretability and usability of connectomic data sets (Bates and Jefferis, 2022).

### 3.1 Machine learning reveals structure-function relationship

We presented a classifier which is able to predict the transmitter identity of a synapse from a local 3D EM volume with high accuracy. We showed that the method generalizes across neurons, brain regions and hemilineages. We started from 34,0000 manually annotated presynaptic locations from 3025 neurons of the Catmaid data set and 1,300,000 automatically detected presynaptic locations from 5902 neurons of the HemiBrain data set. For these neurons we had positive results for their transmitter expression based on immunohistochemical or RNA sequencing results reported in the literature, across 21 different studies. Despite the different image contrast and resolution, we were able to train our model and apply it to tens of millions of automatically detected presynapses across both the FAFB and HemiBrain data sets.

Importantly, we show that individual neurons (Fig. 4c) and cell types (Fig. 11b) that have been matched across the two different brain data sets, FlyWire and HemiBrain, largely concur in their transmitter prediction (92.3%). Individual neurons (Fig. 11a) and cell types (Fig. 11b) that are less confidently predicted in one data set, also tend to have a lower confidence in the second data set (Fig. 11a,b). We also show on a large test set that the classifier generalizes across neurons from different developmental units (*i*.*e*., that derive from different hemilineages) and neuropils, indicating that the influence of the transmitter on the phenotype of a synaptic site is largely conserved across cells of different developmental origins and in different regions. Together this suggests that the key factors that can confuse our network are biological in origin (*e*.*g*., perhaps co-packaging or co-expression of transmitters), rather than data set specific issues.

Given that the relation of synaptic phenotype and transmitter identity is not fully understood in *D. melanogaster*, our model provides an exciting opportunity to learn about the features the classifier uses to predict transmitter identity. However, understanding machine learning models is a challenging problem, especially for contemporary deep learning models due to their intrinsic nonlinearities. Explaining what features and rules a deep learning classifier uses to make its predictions is, in fact, the topic of an active field of research (so-called “explainable AI” or XAI). We build a novel XAI method on top of existing gradient-based methods to highlight class-relevant differences between images of different transmitters (Eckstein et al., 2023). Crucially, our method creates counterfactual synthetic images to investigate class differences. Those counterfactuals allow us to not just highlight areas of importance, but also to show *how* features of a synapse change, if it was of another transmitter. We could use this method to manually identify at least one discriminating feature between each pair of transmitters. A manual validation of these features between the classical transmitters GABA, acetylcholine, and glutamate confirmed our predictions with very high confidence.

We expect our pipeline to be transferable to other connectomic data sets from other species. Because humans cannot tell transmitters apart in *D. melanogaster, D. melanogaster* represented a hard case for transmitter prediction, especially for distinguishing acetylcholine, glutamate and GABA. Since in vertebrate EM data sets, human annotators are able to note differences in the vesicles for excitatory and inhibitory synapses (Gray, 1959; Colonnier, 1968; Tao et al., 2018; Atwood et al., 1972; Uchizono, 1965; Svara et al., 2022), the challenge in vertebrates is likely easier. Symmetric vesicles (usually inhibitory) have already been disambiguated from asymmetric ones (usually excitatory) automatically and at scale (Dorkenwald et al., 2017). Analogous work has also been performed in another invertebrate, the *Ciona intestinalis* larva, with far fewer neurons. Zhang et al. (2022)detected synapses and whether synapses were excitatory or inhibitory. They were able to add signs for 49 neurons with previously unknown transmitter expression. In contrast, our method has shown that it is possible to directly predict transmitter use at synapses across the six most commonly studied transmitters. Since there are many transmitter types, knowing which is expressed is more powerful than delineating only negative from positive. Indeed, glutamate in the insect could play either role differently from acetylcholine and GABA. Because the insect is a harder case for this classification task, we suspect that using our methods in vertebrate connectomic data sets will yield performant predictions with less ground truth required.

### 3.2 Neuronal cell types and hemilineages have consistent transmitter usage that is stereotyped between brains

Our transmitter predictions have confirmed expected, but so far unproven, expression rules in *D. melanogaster*. The neuron-level transmitter predictions across the axon and dendrite in the same neuron (Fig. 7d), across members of a cell type in the same brain hemisphere (Fig. 4d), across neurons matched between hemispheres (Fig. 4b), neurons matched between two separate fly brains (Fig. 4c) and within a hemilineage (Fig. 13a), appear to be consistent. In cases where it is not, our prediction confidences are significantly lower (Fig. 4d-f). This may mean that our results have been inaccurate for a biological reason.

We further find that Lacin’s law largely holds, except in a few cases where mixed expression mainly divides along gross morphological lines (Fig. 6c). We know that as a hemilineage develops, there is a progression in the morphologies that it generates (Lee et al., 2020). It is possible that during this progression, it may switch from producing neurons of one transmitter usage, to neurons of another (e.g. Fig. 6e-h). This would result in cohesive morphological super groups before and after each switch. We might therefore formalize Lacin’s law as: *“Neurons that develop together, as a hemilineage or cohesive sub-division of a hemilineage, share their transmitter usage”*. In a few cases, hemilineage transmitter mixes do not seem to break down along obvious morphological lines. The most common mix is glutamate and GABA (Fig. 6c). Some neurons are known to express both (Das et al., 2011), potentially as much as 3% of the brain (Croset et al., 2018), which could contribute to this issue. Another potential source of error is the mis-assignment of neurons to hemilineages.

Our hemilineage-level transmitter predictions have improved the usability of the transmitter predictions we have made in *D. melanogaster*. Not all neuron-level transmitter prediction are high confidence (Fig. 5b). By taking the assumption that neurons in a hemilineage all express one of acetylcholine, glutamate or GABA, we could assign all members of the hemilineage the same neuron-level transmitter prediction, effectively reducing our small proportion of incorrect neuron-level transmitter predictions. Assuming that Lacin’s Law largely holds, investigators need only assign their neurons to a hemilineage (or morphological super group) to know which fast-acting transmitter a neuron uses (with a few caveats, see Fig. 6b,e-f). This could work for as many as ∼90% of the brain (most “secondary” neurons). This will be easier than matching neurons to their reported cell types (Bates et al., 2019) and easier than again directly predicting transmitter identity from synapse EM data, as we have done here.

### 3.3 The first large-scale survey of neurons of different transmitter expression classes

Our method scales to entire connectomes.

We generated synapse-level transmitter predictions for all synapses in the HemiBrain (9,496,607) and FlyWire (130,107,424) data sets. We amalgamated these into neuron-level transmitter predictions for 11,277 un-truncated neurons in the HemiBrain data set, and 27,706 well-reconstructed neurons from FlyWire of an estimated ∼32,500 central brain neurons (Schlegel et al. in prep). A whole brain single-cell sequencing atlas estimates that 45% of brain neurons are cholinergic, 10% GABAergic, 15% glutamatergic (Davie et al., 2018), which are comparable to our FlyWire results (Fig. 5a).

A new study in the fly visual system has demonstrated, in contrast to longstanding neuroscientific opinion, that it is possible to simulate connectomes directly, correctly predicting neural responses to dynamic stimuli (Lappalainen et al., 2023). However this depended on combining both synaptic weights from the connectome and signs inferred from neurotransmitter expression. This was possible because the repeated columnar architecture of the fly optic lobe contains around 40,000 cells but only around 250 intrinsic cell types, for most of which neurotransmitter identity was either known (Davis et al., 2020) or could be guessed by experts. Our brainwide predictions for ¿5000 cell types provide the key missing ingredient that could allow these simulation methods to be applied at whole brain scale.

We have found axo-axonic connections, particularly where the source axon is cholinergic, to be very common. While about a third of connectivity in the brain is accounted for by cholinergic axons targeting dendrites, we were surprised to find that a high proportion of the brain’s synaptic budget is spent on axo-axonic connections, about a fifth. Inhibitory connections accounted for two thirds of the axo-axonic budget. However, on an individual neuron level there is great diversity in input profiles, in terms of the transmitter types received and the compartments that receive them (Fig. 8c). Axons can often receive a skewed excitatory:inhibitory connection ratio (Fig. 7e). This was less common with dendrites for which the proportion of excitatory input is usually greater (Fig. 8e,f).

It is interesting to consider the role of glutamatergic neurons, versus cholinergic or GABAergic cells. Cholinergic action is known to excite downstream partners (Bossy et al., 1988; Schuster et al., 1993). GABAergic inhibition has been implicated in “divisive inhibition” (Wilson and Laurent, 2005a; Olsen and Wilson, 2008; Olsen et al., 2010). Compared with glutamateigc neurons, GABAerigc neurons have more upstream and downstream synaptic partners (Fig. 8c). What is the role of glutamate in the nervous system? Not enough work has been done to demonstrate its impact in neural circuits across the brain. Its action on downstream partners depends on the glutamate receptor expression in those neurons. At the neuromuscular junction in the peripheral nervous system it is known to have an excitatory action (Jan and Jan, 1976), while neurons in the olfactory system, central complex, superior brain and visual system are inhibited via GluCl channels (Liu and Wilson, 2013; Lu et al., 2022; McCarthy et al., 2011; Molina-Obando et al., 2019). Recent work in the visual system has shown that coincident of cholinergic excitation and cessation of glutamatergic inhibition onto the same neuron can result in a non-linear excitatory effect, a “multiplicative dis-inhibition” (Groschner et al., 2022).

Glutamatergic and cholinergic neurons share many similarities. They tend to be more alike in size (Fig. 14a), may have a similar cellular metabolism (Fig. 5c), are more similarly polarised in their axon-dendrite split, more often project their axons away from their dendrites (Fig. 7c) and tend to target neurons in deeper layers (Fig. 9c). In other ways, they are more like GABAergic neurons.

There are similar numbers of both in the brain (Fig. 5a), their inputs reach targets in similar proportion (Fig. 5b) (Fig. 7) and they fluctuate through brain layers similarly (Fig. 9c). In addition, in the few hemilineages in which a large proportion of two transmitter types seem to intermingle without a clear morphological divide, those transmitters are usually glutamate and GABA (Fig. 6d).

It seems likely that glutamate plays a key inhibitory role across at least the brain. However, the roles of GABA and glutamate input to the same neuron may still have difference effects. Indeed, it has been shown for a set of circadian neurons that applying GABA does not abolish synchronous membrane fluctuations but applying glutamate does (McCarthy et al., 2011), and that glutamate but not GABA is involved in coincidence detection in the antennal lobe (Das et al., 2011). Future work will help uncover the role of glutamate and how its action differs from both acetylcholine and GABA in the *D. melanogaster* brain, hopefully with guidance from the resources we have produced.

Our method, and the data we have made available for *D. melanogaster*, could be used to discover new neuromodulatory neurons in the brain (Fig. 12). It was beyond the scope of this study to validate predicted expression of dopamine, serotonin and octopamine in unexpected or novel neurons (Fig. 12), but we hope the community may be able to use this information as a start point for exploring neuromodulation in the fly central nervous system.

### 3.4 Limitations of the current method

A likely source of transmitter miss-classification is the possibility that a given neuron releases more than one transmitter at each of its synaptic sites. RNA sequencing work (Croset et al., 2018) has shown a preference for some neuropeptides and monoamines to co-transmit with particular fast-acting transmitters. For example, sNPF, CCHamide-2, Tk, space blanket, jelly belly and amnesiac are commonly expressed in cholinergic neurons, Diuretic hormone 31 is commonly expressed in GABAergic neurons and NPF, Neuropeptide-like precursor 1 and Allatostatin A are commonly expressed in glutamatergic neurons. Indeed, in mammals some cases of co-packaging of fast-acting clear core vesicular transmitters are known, including transmitters of opposing downstream effect (Kim et al., 2022). In the fly antennal lobe, some local neurons also seem to co-transmit both glutamate and GABA (Das et al., 2011).

Fortunately, single cell transcriptomic data of the *D. melanogaster* brain shows that transmitter gene expression is largely exclusive between the fast-acting transmitters acetylcholine, GABA and glutamate (Davie et al., 2018; Croset et al., 2018). Although Croset et al. (2018) found that as much as 16% of brain neurons may co-express a fast-acting transmitter (notwithstanding multiple cell captures in the RNA sequencing), it should be noted that a neuron that *co-expresses* transcripts related to multiple fast-acting transmitters usually only *co-transmit* one, *i*.*e*. only one is used at the synapse (Lacin et al., 2019). However, there is widespread co-expression of fast-acting transmitters with other transmitters. For the considered monoamines (serotonin, octopamine and dopamine), co-expression with a fast-acting transmitter is more probable. In particular Croset et al. (2018) suggests that a large fraction of octopaminergic neurons likely co-express glutamate, while serotonin and dopamine show less evidence for co-expression with fast-acting transmitters.

If in some cases, different presynapses across the same neuron transmitted different transmitters from one another, it is possible our network could distinguish them. However, co-transmission most likely occurs as either independently packaged vesicles “coreleased” presynaptic sites (transmitter A and B packaged in distinct vesicles) or as “co-packed” vesicles (A and B together in the same vesicles) (Kim et al., 2022). Therefore, if a particular neuron in the data set were to co-release a fast-acting transmitter and a monoamine the presented classifier would predict only one of the two. In either case, the different kinds of co-transmission presynapses would need to be trained on as new classes for our classifier. However, due to a lack of known and annotated neurons with co-transmission of the considered transmitters in *D. melanogaster*, our current model ignores this possibility (though note that all known octopaminergic neurons in the fly brain are also glutamatergic (Sherer et al., 2020; Croset et al., 2018),so our predicted octopaminergic neurons are also glutamatergic).

Another key limitation is the fact that we only consider the set of six transmitters (GABA, acetylcholine, glutamate, dopamine, octopamine, serotonin). Moreover, due to our use of a softmax normalization at the network output layer, the model is forced to select one of the six classes, even if there is no evidence for any of them. As a result, the current model is not able to identify synapses or neurons that are not part of the considered transmitters, note-ably histamine (Nässel, 1999), tyramine and a vast number of neuropeptides (Croset et al., 2018). Similar to co-expression, we expect an extension to further transmitters to be possible with available training data.

### 3.5 The significance of automatically assigning biological features

Detecting other biological structures that can be seen in EM data could help us further annotate the connectome with features that are informative about neuronal function. For example, a key determinant of how a downstream neuron responds to transmitter release are the transmitter receptors it expresses on its postsynaptic membrane. If it were possible to predict the postsynaptic composition from EM data, (there is some hope (Rickgauer et al., 2017)) we could improve sign labelling in the connectome significantly. As of yet, publicly available ground truth data for this feature is insufficient in *D. melanogaster*. Another example is the detection of mitochondria (upcoming publication in the HemiBrain data set, pers. comm. S. Plaza, data already available Plaza et al. (2022)), which is perhaps indicative of a neurons’ average energy consumption. The detection of individual vesicles, especially dense core vesicles, would help indicate which neurons express neuropeptides and other less well studied transmitters, and may help us get to grips with mixed use synapses. Detecting presynaptic vesicles and postsynaptic density sizes could help understand the relative weight of individual synaptic connections. Detecting likely sites for gap junctions would help us understand which neurons may be directly electrically coupled. Detecting microtubular cytoskeleton could assist neuronal proofreading and assist in further compartmentalising neurons into their ‘trunk versus twig’ structure (Schneider-Mizell et al., 2016), perhaps providing a means by which to weight synapses. In addition, meta data on cell type and developmental lineage will help us break down neurons into genetically related groups that co-vary in important neuronal traits, for example, as we have shown here, transmitter expression.

More generally, our study demonstrates that it is possible both to predict low level cellular properties including key molecular descriptors of cellular function directly from EM data using high-level labels *and* to discover the specific EM features used for these prediction. In addition to details of the network architecture and training, the success of our approach likely depended on three properties of the data that we selected.

First, for both training and inference we used a specific sub-cellular domain (the synapse) likely to contain image features related to our molecules of interest (transmitter secretion systems and their receptors). Second, we aggregated these features on a per cell basis using the independent segmentation of the cell morphology; this was crucial for linking ground truth labels and input data and also improved prediction accuracy. Third, by using multi-modal matching of neuronal morphologies between EM and light level data we were able to bring external molecular ground truth data to bear enabling us to build an expansive ground truth data set.

We believe that variations on this approach have widespread potential in the biomedical sciences. For example, high resolution EM imaging has recently identified new sub-cellular structures (Xu et al., 2021). Our methodology could be repurposed to identify differences in sub-cellular structures associated with discrete cell types in a range of non-neuronal tissues. Similarly it may be possible to identify states including those associated with pathologies.

## 4 Methods

### 4.1 Electron micrograph data sets

The HemiBrain connectome (Scheffer et al., 2020a) has been largely automatically reconstructed using flood-filling networks (Januszewski et al., 2018) from data acquired by focused ion-beam milling scanning EM (FIBSEM) (Knott et al., 2008) and automatically annotated with synapses, followed by manual intervention. Pre- (T-bars) and postsynapses were identified completely automatically. The data can be accessed via the NeuPrint connectome analysis service (Plaza et al., 2022). Automatic segmentation works well in FIBSEM data sets such as the HemiBrain, both because the image-registration required after an acquisition is minimal and the Z resolution achieved is typically 5x superior, as compared with serial section transmission EM (ssTEM). Unlike the HemiBrain, the FAFB ssTEM image data comprises an entire female fly brain. Two autosegmentations of the data exist (Li et al., 2020b; Dorkenwald et al., 2022), we used Dorkenwald et al. (2022) and automatically detected synapses (Buhmann et al., 2019) for our biological analyses. However, to build out ground truth data we used high-fidelity manually reconstructed neurons and synapses built using CATMAID (Schneider-Mizell et al., 2016). Because the HemiBrain is a *dense* connectome where a complete attempt (with an associated failures rate) has been made to detect every neuron and synapse, we can use the HemiBrain data set for certain types of analysis where we cannot use the fafb-catmaid or FlyWire data set. The fafb data sets, at the time of writing, is a sparsely reconstructed data set where there are some systematic gaps in the data. It is expected to be a dense connectome within the year. However, at present, it would be inaccurate to, say, sum the number of synapses for each transmitter prediction in each neuropil in FAFB because some neurons/neuronal arbor in each region are still missing.

### 4.2 Neuronal reconstructions

We excluded neurons that were truncated so as to loose a large portion of their axon or dendrite in this data set, for example brain descending neurons, neurons with fewer than 100 presynapses. For biological analyses (figures 5-9) we also excluded Kenyon Cells and neurons that had not previously been identified as octopaminergic (Busch et al., 2009) because we had a high confidence that these predictions were incorrect. In so doing we selected 11,277 from the HemiBrain data set. These neurons were semi-automatically reconstructed by Scheffer et al. (2020a). For the FlyWire data set we used 27,706 intrinsic brain neurons. These neurons were selected because they had been well reconstructed in FlyWire by the end of 2022, and we could “skeletonize” and “split” into a separable axons and dendrites (Bates et al., 2020a; Schneider-Mizell et al., 2016) for our biological analyses. They were built by 1,366,543 edits of automatically reconstructed segments (Dorkenwald et al., 2022), from 100 human annotators (see Acknowledgements). Our synapse-level transmitter predictions can be associated with these segments and therefore neurons built by the community. We have not yet cross-matched our full pool of neurons from both data sets (only 469 neuronal cell types).

#### 4.2.1 Neuron level transmitter predictions

To calculate our neuron-level transmitter predictions, we determined which synapse-level transmitter prediction was most common for each neuron. If the difference between the top and second highest transmitter was < 10% a neuron’s neuron-level transmitter prediction was designated “unknown”. To determine a neuron-level transmitter prediction confidence score for each neuron (or neuronal compartment), we summed the number of predictions per synapse, weighted by the corresponding values in the confusion matrix column for the winning transmitter and normalized the result by the total number of presynapses for each neuron. For example, say we determined a FlyWire neuron was cholinergic because a majority of its pre-synapses were predicted to transmit acetylcholine. Each pre-synapse predicted for acetylcholine would get a score of 0.95 - the proportion of cholinegic ground-truth pre-synapses correctly determined as cholinergic (Fig. 2b). Any pre-synapse presumably mispredicted as GABA in our cholinergic neuron got a score of 0.02 (the proportion of ground-truth cholinergic pre-synapses mispredicted as GABAergic, Fig. 2b), if it was predicted glutamate it got a score of 0.02, etc. The neuron’s final confidence score is the mean of these values.

We removed potential erroneous presynapses by employing a threshold for synapse detection (a cleft score above 50 for FAFB auto-detected synapses, and a confidence score above 0.5 for HemiBrain auto-detected synapses) and presynapse count per neuron (100). Our HemiBrain thresholds were chosen in alignment with the flyEM HemiBrain project (Scheffer et al., 2020b). We validate our cut-off for automatically-detected FAFB presynapses (Buhmann et al., 2019) by evenly sampling 4,306 across all 6 top synapse-level transmitter predictions and neuronal compartments (see Fig. 7a) and determining if the auto-detected preynapses were true synapses using a human annotator. 32% of auto-detected presynapses were erroneous. Our chosen threshold eliminates ∼13% of valid presynapses and ∼60% of erroneous detections (Fig. 10a). Because a higher proportion of presynapses on the primary dendrite and cell body fiber tract were erroneous, we excluded all presynapses in these neuronal compartments as well as in the cell body (Fig. 10b), retaining only axonic and dendritic presynapses. It is notable that a larger proportion of octopamine predicted presynapses prove erroneous (43%) indicating that the network may guess octopamine for dark non-synaptic bodies it is asked to examine (Fig. 12).

**Figure 10:**
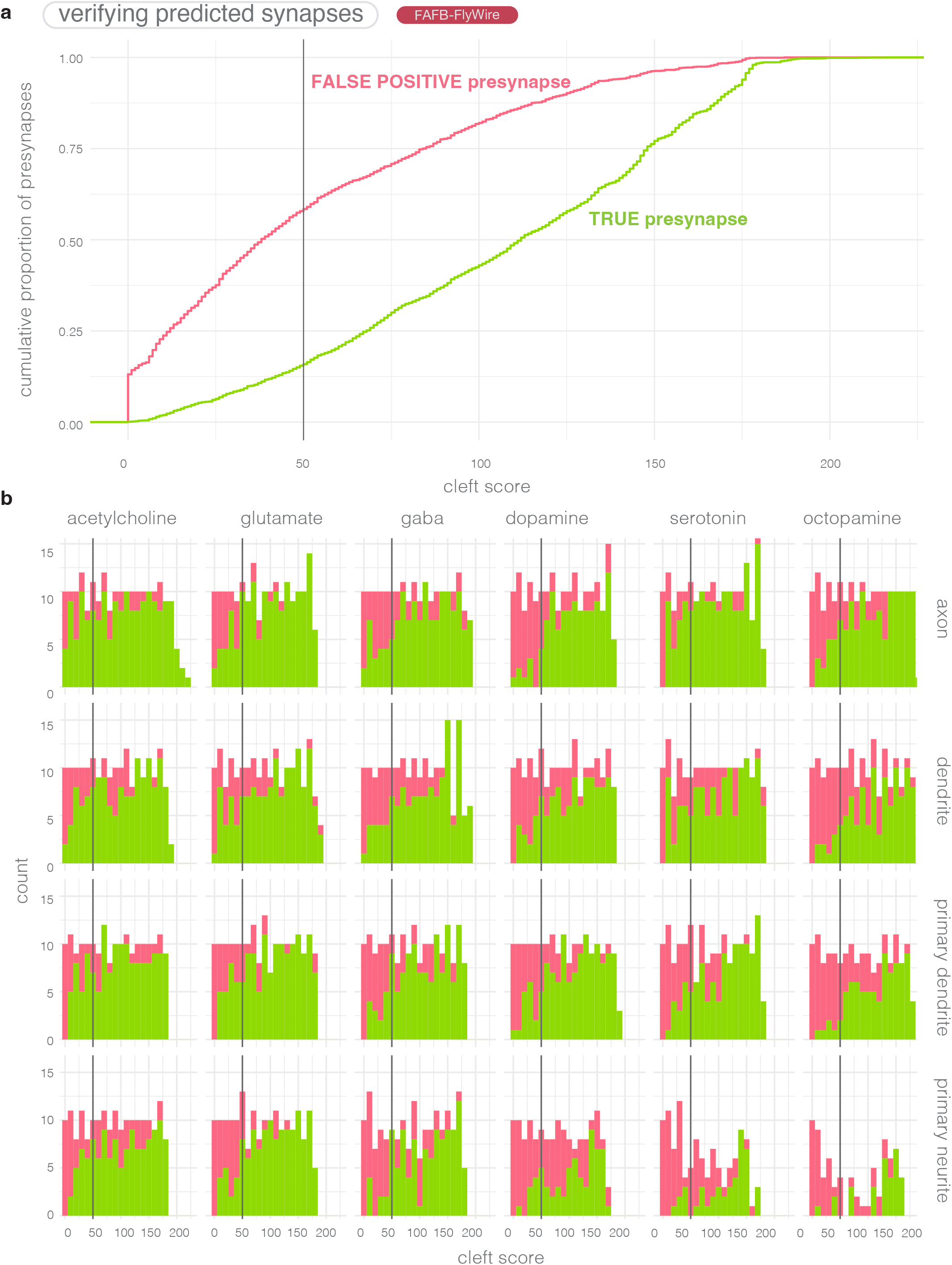
Human review of auto-detected presynapses in the FAFB-FlyWire data set **(a)**: Empirical cumulative density distribution curve for review of 4,306 automatically detected preynapses from the FAFB-FlyWire data set. A presynapse comprises the synaptic machinery and vesicles on the source neuron’s side of the synaptic cleft (they are the output zones of a neuron). Detected preynapse have a “cleft score” that ranges between 0 to over 200, which indicates how discriminable the synaptic cleft at the presynaptic site is for the detection network (Buhmann et al., 2019). Our threshold of 50 indicated as a vertical grey line. Green, determined to be a true presynapse by a human annotator, pink, determined to not be a true presynapse. **(b)**: Rates of false presynapse detection across cleft scores, transmitter types and compartments. We sampled ∼180 for each set of conditions. We sampled ∼10 presynapses per cleft score bin (width 10), presynaptic transmitter prediction type (columns) and neuronal compartment (rows). Histograms show the number of presynapses determined to be real (green) or not (red) by a human annotator (A.S.B). Presynapses were reviewed using the FlyWire interface (Dorkenwald et al., 2022).

**Figure 11:**
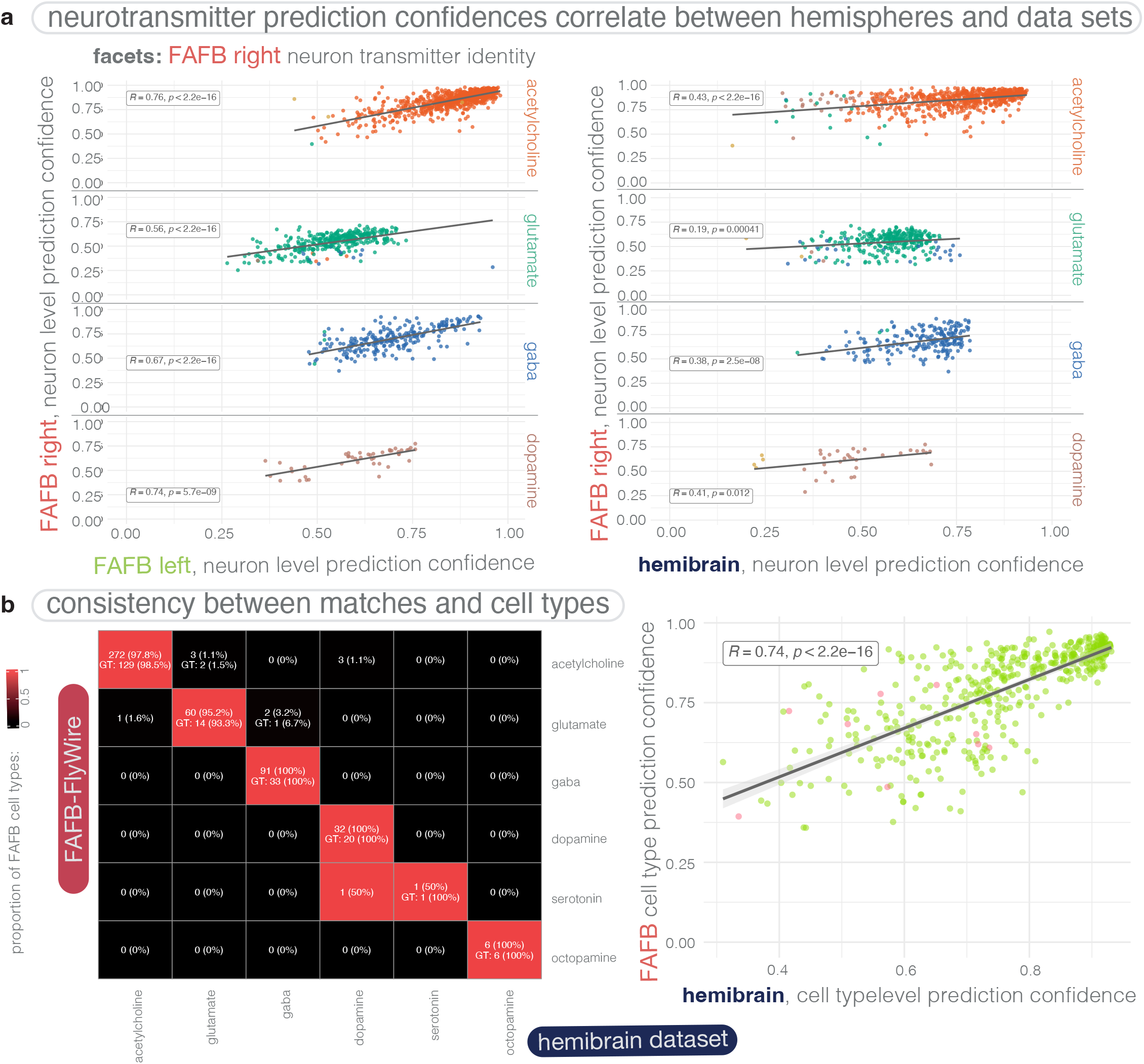
Comparing neurons’ transmitter usage predictions between connectome data sets from separate animals (FAFB-FlyWire and hemibrain) and between two hemispheres in the same data set (FAFB-FlyWire). **(a)**: A scatter plot comparing neuron-level confidence scores in the transmitter prediction of FAFB-FlyWire data set central brain neurons, faceted by the neuron-level transmitter prediction for the FAFB-FlyWire right side homolog. Individual points coloured by their FAFB-FlyWire left side neuron-level transmitter prediction score. There are 878 acetylcholine, 382 glutamate, 272 GABA, 45 dopamine, 18, serotonin, 7 octopamine matches by the FAFB-FlyWire right side prediction, of which 40 (2.5%) disagree with the FAFB-FlyWire left side prediction. **(a)**: A scatter plot comparing neuron-level confidence scores in the transmitter prediction of hemibrain data set central brain neurons. Individual points coloured by their hemibrain side neuron-level transmitter prediction score. There are 630 acetylcholine 329 glutamate, 209 GABA, 39 dopamine, 7, serotonin, 4 octopamine matches by the FAFB-FlyWire right side prediction, of which 94 (7.7%) disagree with the hemibrain prediction. **(c)**: A confusion matrix showing the neuronal cell type level prediction (mode of the neuron-level transmitter predictions per neuronal cell type) for neuronal cell types in the FAFB-FlyWire and hemibrain data sets. Cells give the number of cross-matched neuronal cell types we examined, and the number of those present in the ground truth data for at least one of the two data sets. **(d)**: A scatter plot showing the correlation between our mean prediction confidence scores for FAFB-FlyWire and hemibrain neuronal cell types. Each point is a neuronal cell type identified in both data sets (469). Green points mean that the transmitter prediction agrees between the two data sets, pink points indicate disagreement. Scatter plots display Pearson’s product-moment correlation, giving R, the coefficient and associated p-value.

**Figure 12:**
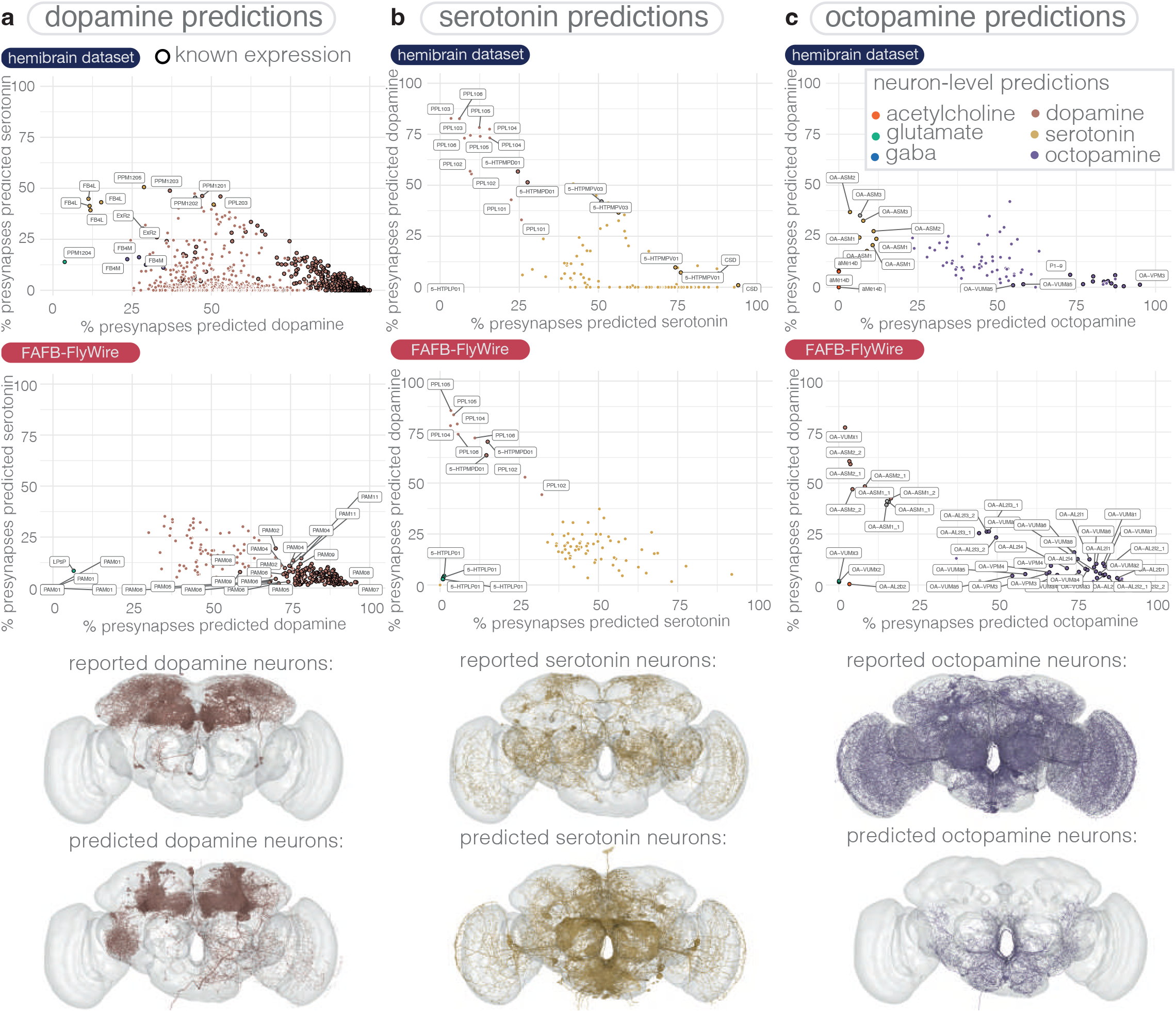
neuron-level transmitter predictions within secondary hemilineages **(a)**: A look at dopamine predicted neurons. We show two scatter plots using data and predictions from the the hemibrain (upper) and FAFB-FlyWire (lower) data sets. The proportion of presynapses in each neuron (each point) that is predicted as dopamine (X-axis) and serotonin (Y-axis) Neurons that have been predicted as dopaminergic, or given as dopaminergic, in the ground truth are shown. Those neurons from the ground truth data are circled with a black ring. Upper brain plot shows neurons known to be dopaminergic (colored by their neuron-level transmitter prediction). Lower brain plot shows neurons predicted to be dopaminergic, excluding those in the upper plot. Many weakly predicted dopaminergic neurons belong to the central complex and mushroom body, where the density of presynapses from other neurons may have contributed to possible mis-predictions (see Methods). Same as *a*, but for serotonin predictions. PPL101-6 neurons are known to co-express dopamine and serotonin but are predicted as dopaminergic. Some known serotonergic neurons have low proportions of presynapses predicted as serotonergic. **(c)**: Same as *a*, but for octopamine predictions. Most octopamine neurons have been identified in prior work. Many of our octopamine predictions (no dark circle) actually indicate neurons that express some other dense core vesicle transmitter in abundance, for example PI neurons which express insulin. Interesting putative octopaminergic aMe14b neurons (Busch et al., 2009) (also known as OA-AL2b2 neurons) are predicted as cholinergic. Busch et al. did note that might not be octopaminergic, as the neurons were not obviously immunoreactive to their probe: “Because not all neurons in cluster AL2 of NP7088 are OA-immunoreactive, and because OA-AL2b2 was identified in NP7088, but not in tdc2-GAL4, this cell type might not be octopaminergic.” OA-ASM neurons also are not predicted octopaminergic, but serotonergic. On them, Busch et al. note: “There are 8 OA-immunoreactive somata localized to the anterior superior medial protocerebrum uniquely labeled by tdc2-GAL4 (the ASM cluster). Yet they are not necessarily octopaminergic, as there are GAL4-positive neurons without OA-immunoreactivity in this cluster.”

**Figure 13:**
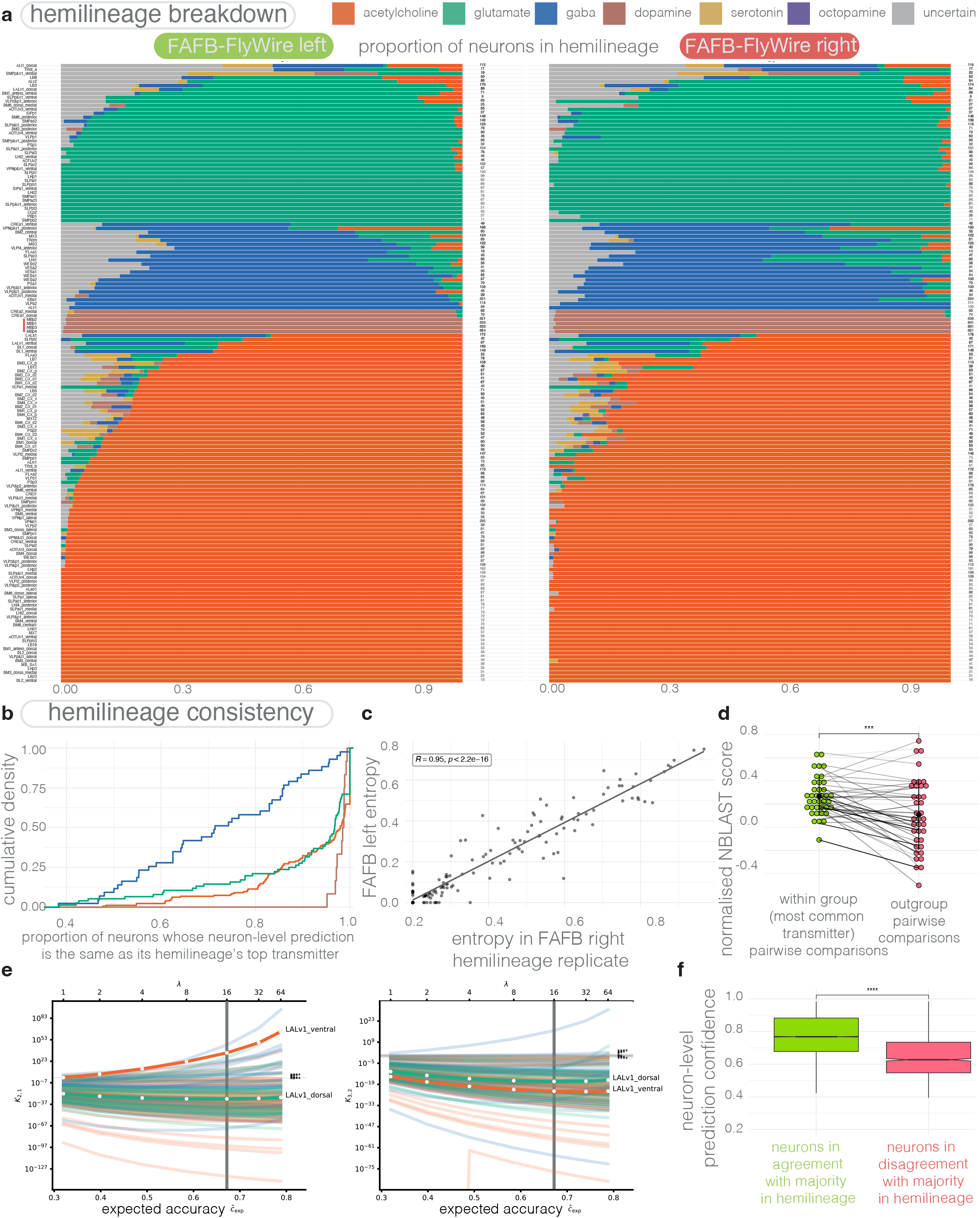
neuron-level transmitter predictions within secondary hemilineages **(a)**: Consistency of neuron-level transmitter predictions within selected hemilineages in the adult *D. melanogaster* brain. Bar plots show the proportion of neurons in each each hemilineage, predicted to express each of our six transmitters. Data shown for neurons of the left hemisphere of the FAFB-FlyWire data set (left), the right hemisphere (right). Hemilineage names are on the left of the bar plots, and the numbers of neurons per hemilineage are on the right. Empirical cumulative density plot shows how consistent transmitter within each hemilineage is predicted to be. The Y axis gives the proportion of hemilineages, and the X axis gives the proportion of neurons in those hemilineages that “voted” for the top transmitter (colour groups). **(c)**: How the Shannon entropy (base 6) in the neuron-level transmitter predictions for each hemilineage correlate, between the hemilineage copy on the right (X-axis) and left (Y-axis) hemispheres of the FAFB-FlyWire data set brain. **(d)**: Dot plot shows the mean normalized pair-wise NBLAST scores (Costa et al., 2016) between neurons expressing the majority transmitter within a hemilineage (each green dot, each member of pair expresses the main transmitter) and between these neurons and those expressing other transmitters (pink), where there are at least 10 neurons that do express another transmitter. Dots represents means taken per hemilineages (161). **(e)**: Bayes factor analysis of likelihood of number of transmitters per hemilineage. The left panel shows the Bayes factor *K*_2,1_, the relative likelihood ratio of two or one transmitter, for a range of rate parameters *λ* and their corresponding expected accuracy. 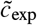 Values greater than 1 indicate two transmitters are more likely, while those less than indicate one transmitter is more likely. The right panel is identical to the left for Bayes factor *K*_3,2_, i.e., values greater than 1 indicate three transmitters are more likely, while those less than indicate two transmitters are more likely. LALv1 hemilineages are highlighted, showing LALv1 dorsal likely has one transmitter, while LALv1 ventral is more likely to have two than one or three. Vertical lines indicate *λ* = 16 corresponding to the one-versus-rest Bayes analysis in Fig. 6c. Evidence strength: *: *K* ≥ 10^1/2^; **: *K* ≥ 10^1^; ***: *K* ≥ 10^3/2^; ****: *K* ≥ 10^2^. (Jeffreys, 1998). **(f)**: The neuron-level transmitter prediction confidences for neurons in agreement with their hemilineage, and those that are not. Box plots show the median (horizontal line), the lower and upper hinges correspond to the first and third quartiles (the 25^th^ and 75^th^ percentiles). Whiskers extend to 1.5 times the inter-quartile range (IQR). Data compared using Wilcoxon two-sample tests. Significance values: ns: p > 0.05; *: p <=0.05; **: p <= 0.01; ***: p <=0.001; ****: p <= 0.0001.

**Figure 14:**
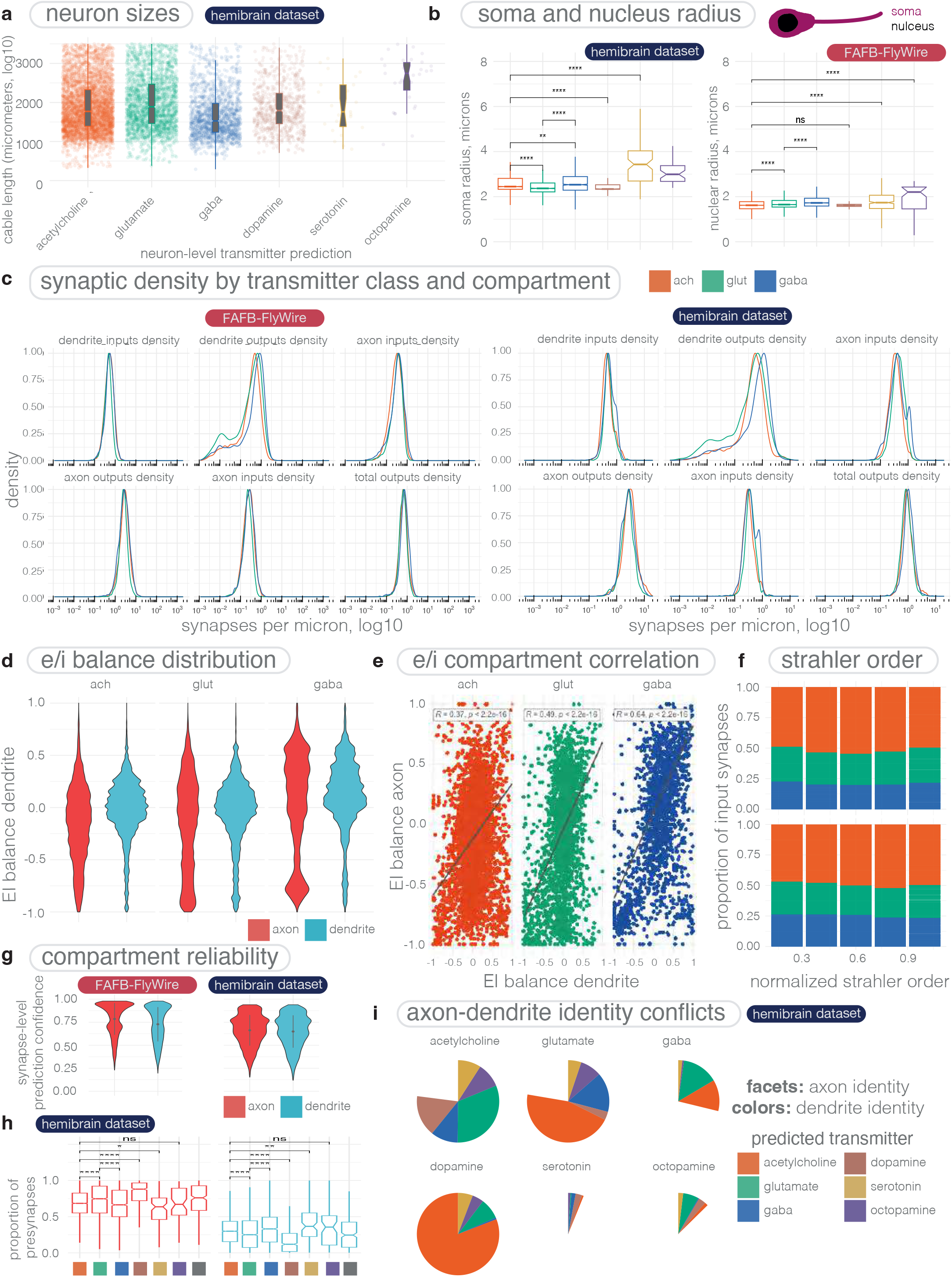
**a:** Cable length by neuron-level transmitter prediction. **(b)**: Soma, *i*.*e*. neuronal cell body, and nucleus size by neuron-level transmitter predictions. The hemibrain data set provides a soma segmentation (left), and the FAFB-FlyWire data set provides a nucleus segmentation (right) (Plaza et al., 2022; Dorkenwald et al., 2022). **(c)**: The distribution of synaptic densities across by neuron-level transmitter prediction and compartment. Y axis, max-normalised proportion of neurons. X-axis, either postsynapses (inputs) or presynapses (outputs) per micrometers of axonic or dendritic cable, transformed by log10. Left, FAFB-FlyWire data set, right, hemibrain data set. **(d)**: Violin plots of EI balance by neuron-level transmitter prediction and compartment **(e)**: Correlation in EI balance between axon and dendrite. Each dot is one neuron. **(f)**: Breakdown of neuron-level transmitter prediction for synaptic inputs by normalised Strahler order. Strahler order is a measure of branching complexity. High Strahler order branches are more central to a neuron’s tree. Lower order ones more peripheral, such that branches with leaf nodes are Strahler order one. The higher Strahler order between neurons can very greatly. Here, the Strahler order has been max normalised by each neuron’s highest Strahler order and binned into four groups, bin width 0.2. **(g)**: The synapse-level transmitter prediction score by compartment. Left FAFB-FlyWire data set, right hemibrain data set. **(h)**: The proportion of presynapses (synaptic outputs) in the axon or dendrite. Boxplot plots shown by pre-synaptic neuron-level transmitter prediction and compartment. **(i)**: Break down of hemibrain axon-dendrite compartment-level transmitter prediction mismatches. Facets indicate the axons’ strongest prediction, and colours the dendrites. The pie charts are proportional to one another, and so can be directly compared. Boxplots show the median (horizontal line), the lower and upper hinges correspond to the first and third quartiles (the 25^th^ and 75^th^ percentiles). Whiskers extend to 1.5 times the inter-quartile range. Significance values: ns: p > 0.05; *: p <=0.05; **: p <= 0.01; ***: p <=0.001; ****: p <= 0.0001.

**Figure 15:**
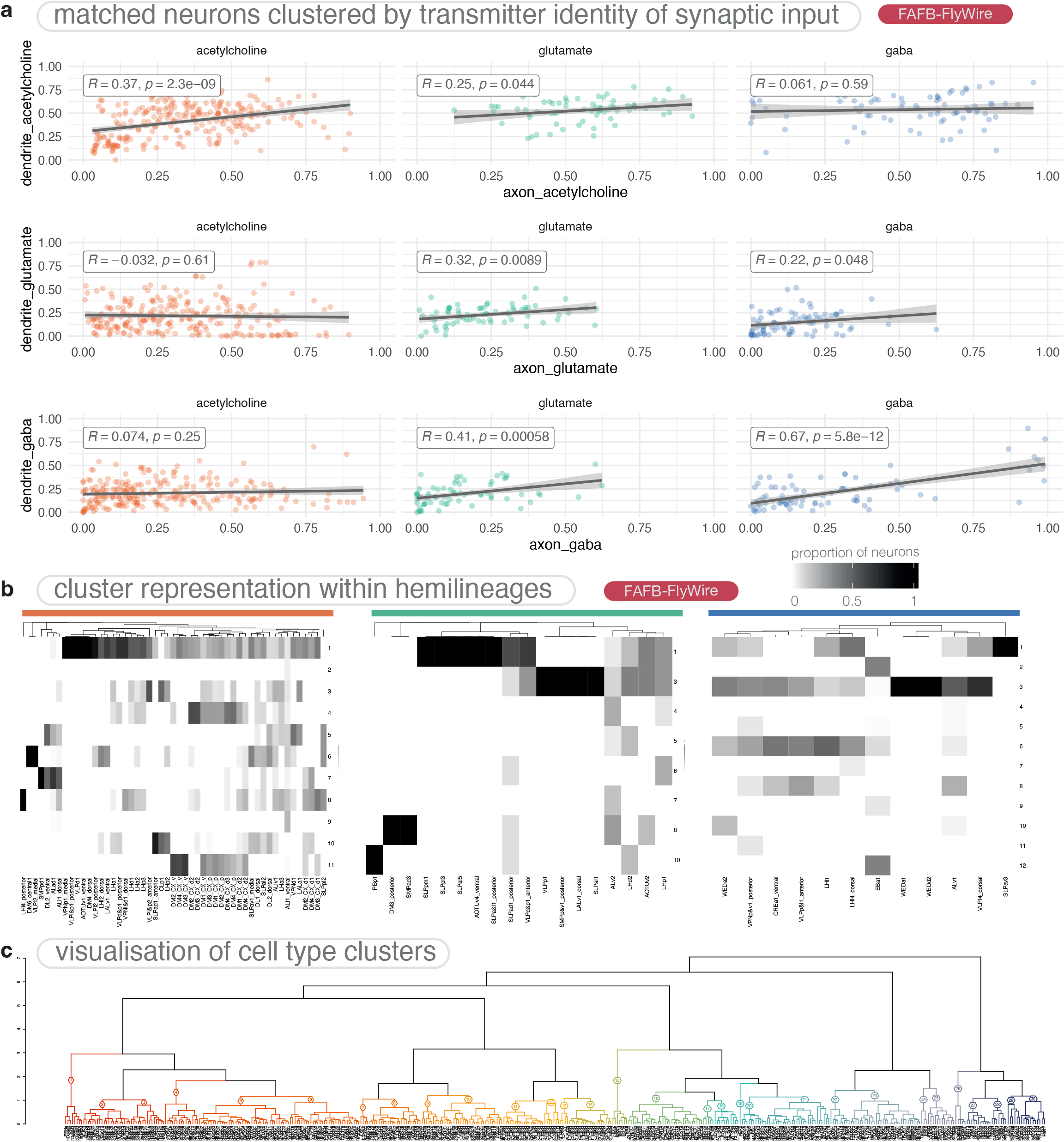
**(a)**: Correlations between opposing input transmitter types by compartment. Plots faceted by the source (upstream) neurons’ neuron-level transmitter prediction (axis values). Colored by the target (downstream) neurons’ neuron-level transmitter prediction. The X axis shows the proportion of a neuron’s input accounted for by the input type on the axis label. Each dot is one neuron. For calculating the R^2^ and p-values, neurons for which a proportion on either the X or Y axis fell below 0.1 or above 0.9 were excluded, to remove outlier cases with a very strong input preference. **(b)**: Cluster representation within FAFB-FlyWire hemilineages. Rows, the clusters shown in Fig. 8a, and here as neuron plots in C. Columns, selected FAFB-FlyWire hemilineages. Plots faceted by majority neuron-level transmitter prediction. Colour scale indicates the proportion of neurons within the hemilineage that belong to each cluster. **(c)**: Dendrogram used for both heatmaps in Fig. 8a, with cell type labels. Coloured by cluster, K = 30.

One caveat is potential image contamination by unwanted presynapses. The cube of image (edge length 640 nm) centered on each predicted presynaptic location and used for prediction was not masked with neuronal identity. It is possible that features from proximal presynapses have skewed the result. This may also be the case for a few neuronal classes, namely Kenyon cells (KCs) of the mushroom body (KCs), olfactory sensory neurons of the antennal lobe (OSNs), the protocerebral bridge neuron LPsP and central complex (CX) neurons such the hDelta, vDelta and FB tangential classes. The mushroom body, central complex and antennal lobe are especially synapse-dense regions of the brain that appear prominently in Nc82 immunohisochemsity synapse stains of the brain. KCs are potentially mis-predicted as dopaminergic, and are in a region of the brain (mushroom body) with a high density of true dopaminergic PPL1 and PAM neurons, which were also represented in our ground truth (KCs were not). Some OSNs are potentially mis-predicted as serotonergic, and are in a region of the brain (antennal lobe) with a high density of true serotonergic presynapses from the CSD neuron, which were also represented in our ground truth (OSNs were not). LPsp are potentially mis-predicted as glutamatergic, and are in a region of the brain (protocerebral bridge) with a high density of true glutamatergic presynapses from the delta7 neurons, which were also represented in our ground truth (LPsp also was). Central complex (CX) hDelta and vDelta neurons (not in our ground truth data) are mostly predicted as cholinergic but some a potentially erroneously predicted as either dopaminergic or serotonergic. They are present in a structure that contains diverse transmitter input, e.g. dopaminergic input from FB7B neurons and dopamine/serotonin input from ExR neurons, and has very dense synaptic connectivity. See (Fig. 5d) for the regional biases.

#### 4.2.2 Hemilineage assignments in D. melanogaster

Our work assigning neurons in the FlyWire data set to hemi-lineages will be reported in an upcoming manuscript (Schlegel et al. 2023, in prep). We briefly detail the process here. Cell body fiber tracts for identified hemilineages had previously been identified using TrakEM2 (Cardona et al., 2012) in a light-level atlas for a *D. melanogaster* brain, stained with an antibody against neurotactin (BP104) (Lovick et al., 2013). We extracted these expertly identified tracts and registered them into a common template brain, JFRC2, using CMTK (Rohlfing and Maurer, 2003), and then into FAFB space (Bates et al., 2020a). We could now identify cell body fiber tracts in this ssTEM data set in the vicinity of the transformed hemilineage tracts using the flywire.ai Web interface (Dorkenwald et al., 2022). We compared our candidate neurons to registered images of lineage clones, produced using genetic tools and light microscopy (Ito et al., 2013; Yu et al., 2013). Hemilineages can be told apart in an image of a lineage clone, by the different placement of their cell body clusters, and the tract their cell body fibers take into the neuropil. In some cases, one of a lineage’s hemilineage may apoptose (Jiang and Reichert, 2012), leaving only a single hemilineage. From light level data that labels a whole lineage and all of its hemilineages together (Ito et al., 2013; Yu et al., 2013; Wong et al., 2013; Lovick et al., 2013), it is occasionally difficult to assess whether there is a single hemilineage or multiple, if the cell body clusters happen to not separate sufficiently. While they can be told apart using developmental genetic tools, e.g. in the case of LALv1 (Lee et al., 2020), We divided the following lineages, which had not already been divided in foundational work (Ito et al., 2013; Yu et al., 2013; Wong et al., 2013; Lovick et al., 2013), into separate hemilineages based on careful assessment of their cell body fiber tracts, neuropil entry sites and first branch point positions: ALl1, DL1, DL2, LHl2, LHl4, SIPa1, SLPav1, VLPl2, VLPl4. The majority of lineages we analyse had already been reported (Ito et al., 2013; Yu et al., 2013; Wong et al., 2013; Lovick et al., 2013), however we discovered five new hemilineages (their pairings into lineages are not known) that we named: LHp3, CREl1, LALa1, SMPpm1, SLPpm2 in the nomenclature of Ito et al. (2013) and Yu et al. (2013), and CP5, DALv3, BAlp2m, DPMm2m and DPLc2 in the nomenclature of Wong et al. (2013) and Lovick et al. (2013).

We have used only the best 161 hemilineages, human reviewed on both the right and left FlyWire hemispheres, in this work. However, assigning neurons to hemilineages is difficult. For example, neurons from the lineage LALv1 co-bundle between its two hemilineages, making their disambiguation dependent on examining light level data and single neuron clones (Lee et al., 2020). Without work from Lee et al. (2020) on this lineage’s detailed composition, we might have incorrectly assumed that its glutamatergic and cholinergic neurons from its two separate hemilineages intermingled in a single hemilineage. High quality light-level data is not available for every hemilineage, meaning that we had to make some judgement calls in our hemilineage discrimination.

#### 4.2.3 Hemilineage transmitter predictions

We retrain the classifier on 90% of the entire data set and use the remaining 10% to select the best performing iteration. We predict the transmitter identity of 42,000,000 synapses within 23,755 neurons with so far unknown transmitter identity. These neurons come from a total of 167 hemilineages. In the following, we analyse the results by quantifying how transmitter predictions are distributed over neurons and synapses within hemilineages.

##### 4.2.3.1 Neuron Level Entropy

In order to quantify multi-modality of transmitter predictions on neuron level within a hemi-lineage we calculate the entropy *H* of the transmitter distribution over neurons in the following way: Let *n* ∈ *N*_*h*_ be a neuron in hemilineage *h* and *ŷ*_*n*_ ∈ *Y* = {GABA, acetylcholine, glutamate, serotonin, octopamine, dopamine} the predicted transmitter of neuron *n*. Then

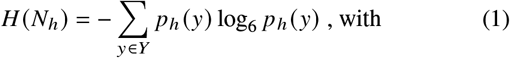

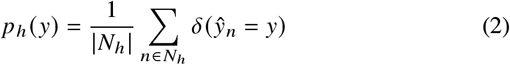

A value of *H* (*N*_*h*_) = 0 then means that all neurons within hemilineage h have the same predicted transmitter, while a value of *H* (*N*_*h*_) = 1 means that within hemilineage h all predicted transmitters are equally common.

##### 4.2.3.2 Synapse Level Entropy

Similarly we can quantify the average multimodality over synapses within neurons of a given hemilineage: Let *s* ∈ *S*_*n*_ be the synapses in neuron *n* ∈ *N*_*h*_ of hemilineage *h* and *ŷ*_*s*_ the predicted transmitter. The entropy of predicted synaptic transmitters *H*(*s*_*n*_) in neuron n is then given by:

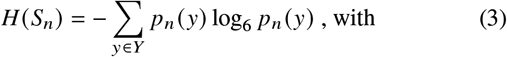

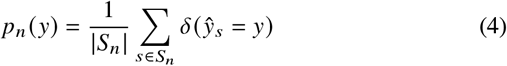

With this the average synaptic entropy over all neurons within hemilineage *h* is given by:

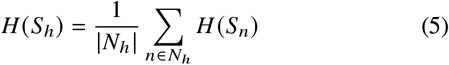

A value of *H* (*S*_*h*_) = 0 means that all synapses of all neurons in hemilineage h have the same predicted transmitter, while a value of *H* (*S*_*h*_) = 1 means that in all neurons within hemilineage h all synaptic transmitter predictions are equally common. Fig. 6d shows the distribution of *H* (*N*_*h*_) and *H* (*S*_*h*_) of all predicted hemilineages with more than ten neurons that have more than 30 synapses each. On the population level we find relatively higher values of *H* (*S*_*h*_) (Synapse level entropy) than *H* (*N*_*h*_) (Neuron level entropy). 75% of hemilineages show a synapse level entropy below *q*_75_ (*H* (*S*_*h*_)) = 0.41 as compared to *q*_75_ (*H* (*N*_*h*_)) = 0.20. This is reassuring as it suggests less variation of transmitter identity of neurons within a hemilineage compared to variations of transmitter identity predictions within individual neurons, meaning there is improved consensus of predictions when aggregating across populations. However, in cases with a high level of synaptic entropy, such as hemilineage TRdl_a, it is less clear whether neuron level multimodality is an artifact of uncertain, multimodal predictions on the synapse level of individual neurons. In contrast, hemilineages such as SMPpd1 show high neuron level entropy *H* (*N*_*h*_) ≥*q*_75_ but low synapse level entropy *H* (*S*_*h*_) ≤ *q*_25_, suggesting clear neuron level segregation of predicted transmitters within those hemilineages. Hemilineages such as ALad1 with (*H S*_*h*_) ≥ *q*_25_ and *H S*_*n*_ < *q*_25_ appear homogeneous within each neuron and within the entire hemilineage.

#### 4.2.4 Probability to observe transmitter predictions ŷ

Given a neuron has true transmitter *y* ∈ *Y*, the probability that we predict transmitter *ŷ* ∈ *Y* (assuming that each prediction is independent and identically distributed) is given by the categorical distribution

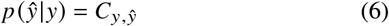

where *C* is the neuron confusion matrix obtained on the test data set (see Fig. 2a).

Let *m* be the number of different transmitters in hemilineage *h*. We model the probability *p* (**ŷ**|*m*) of observing transmitter predic-tions **ŷ** = {*ŷ*_0_, *ŷ*_1_, …, *ŷ*_*n*_} under the assumption that hemilineage *h* contains *m* different transmitters. Here, *ŷ* _*j*_ is the predicted transmitter of neuron *j* in hemilineage *h* with *n* neurons total. Let ℙ_*c*_ (*Y*) be the set of subsets of true transmitters *Y* with cardinality *c*, then:

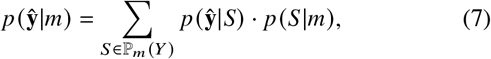

where *p* (**ŷ**|*S*) is the probability to observe predictions **ŷ** if the hemilineage has true underlying transmitters *y* ∈ *S* and *p* (*S*|*m*) is the probability for the set of true transmitters *S* given the hemilineage contains *m* different transmitters. Since we assume i.i.d. predictions **ŷ**, *p* (**ŷ**|*S*) factorizes as follows:

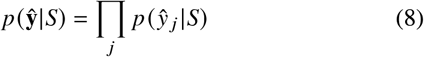

and marginalizing over *y* ∈ *S* yields:

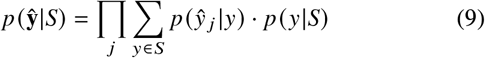

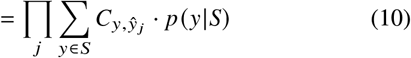

Regarding *p* (*S*|*m*) and *p* (*y*|*S*) we assume a flat prior, i.e.:

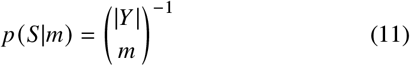

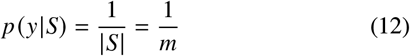

With this, the probability of observing predictions **ŷ** given *m* different transmitters becomes:

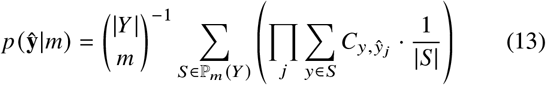

4.2.4.1 Bayes Factor: With this formalism in place, we can compare hypotheses about the number of true transmitters *m* in a given hemilineage by using the Bayes Factor 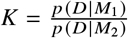, where D is our observed data (predicted transmitters) and *M*_1_, *M*_2_ are two models about the underlying true transmitters that we wish to compare. The Bayes factor for a model *M*_1_ with *m*_1_ true transmitters per hemilineage and model *M*_2_ with *m*_2_ different transmitters is given by:

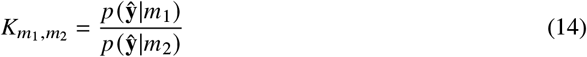

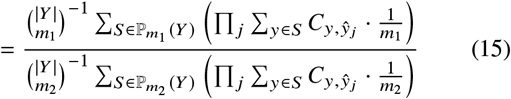

So far, we assumed that 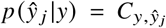, i.e. we estimate this distribution on the test data set. However, because our test set is finite we can not expect that the estimated error rates perfectly transfer to other data sets. In order to relax our assumptions about this distribution we simulate additional errors, by incorporating additive smoothing on the counts of neurons *N*_*y,ŷ*_ that have true transmitter *y* and were predicted as transmitter *ŷ*, i.e.:

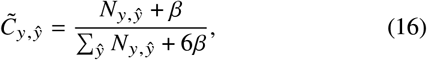

where *β* ∈ ℕ_0_ is the smoothing parameter. With 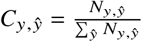 we then have

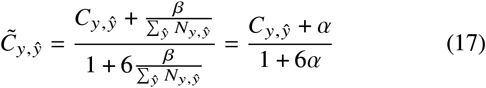

and *α* ∈ ℝ_≥0_ the count normalized smoothing parameter. In the limit of *α* → ∞, 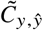 approaches the uniform distribution with probability 1/6 for each transmitter, whereas a value of *α* = 0 means we recover the observed confusion matrix *C*. With this our distributions are now parametrized by *α* and the bayes factor becomes:

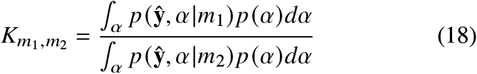

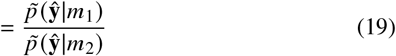

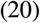

where 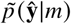 is as defined in (13) but with *C*_*y,ŷ*_ replaced with its expected value 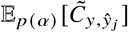.

The prior distribution on *α, p α* allows us to encode our prior knowledge about *α* and use it to weight the likelihood of the corresponding model. Given the data, a value of *α* = *ϵ* with epsilon small (0 < *ϵ* ≪ 1), should be most probable, while the probability of values *α* > *ϵ* should monotonically decrease as we deviate more from the observed confusion matrix. Values of *α* < *ϵ* should have probability zero, because they correspond to the un-smoothed confusion matrix with zero entries, i.e. a probability of zero for mis-classification of certain transmitters. While these probabilities may be small, they are likely greater than zero and an artifact caused by the finite test set. Many distributions fulfill these criteria, in particular the family of exponential distributions with rate parameter *λ*:

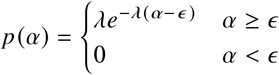

Thus *λ* controls the weight for smoothing parameter *α* in the integral 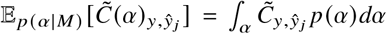. For *λ* → 0, the expected confusion matrix approaches the unweighted average of all *C* (*α*) in the integration range. For *λ* → ∞, the expected confusion matrix approaches the *ϵ*-smoothed confusion matrix 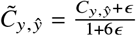.

The rate parameter *λ* can also be understood via its influence on the expected average accuracy 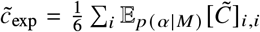. For values of *λ* → 0, the expected accuracy approaches chance level while for values of *λ* → ∞, the expected accuracy approaches the *ϵ*-smoothed, observed accuracy on the test set.

To summarize overall maximum likelihood of number of true transmitters in a given lineage, for a fixed *λ* we consider a one-versus-rest Bayes factor:

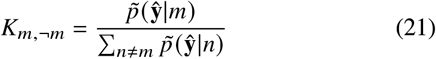

### 4.3 Connectome analysis tools

All software tools used in this study are summarized in Table 2.

**Table 2:**
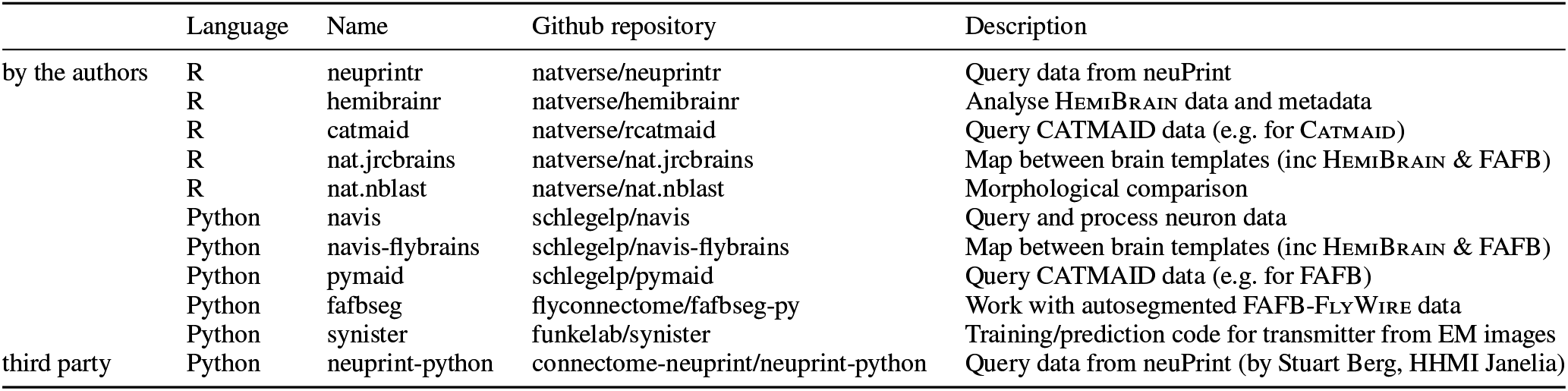
R and Python packages used in this study.

## 5 Acknowledgements

This work was supported by a Wellcome Trust Collaborative Award (203261/Z/16/Z), National Institutes of Health (BRAIN Initiative grant 1RF1MH120679-01), an ERC Consolidator grant (649111), a Neuronex2 award (MC_EX_MR/T046279/1, NSF reference: 2014862), with additional core support from the MRC (MC-U105188491) to G.S.X.E.J; by the Howard Hughes Medical Institute at Janelia Research Campus (J.F.); a Boehringer Ingelheim Fonds PhD Fellowship, EMBO fellowship (ALTF 1258-2020), and a Sir Henry Wellcome Postdoctoral Fellowship (222782/Z/21/Z) to A.S.B.. A.S.B. thanks Rachel I. Wilson for her support and interest as he finished this project after having moved to the Wilson lab. The manually placed FAFB synapses in this study were identified in Catmaid (Zheng et al., 2018; Schneider-Mizell et al., 2016). Catmaid is a collaborative environment in which 27 labs have participated to build connectomes for specific circuits. For these annotations, we thank Ruairi Roberts, Fiona Love, Lisa Marin, Amelia Edmondson-Stait, Xincheng Zhao, Jawaid Ali, Johann Schor, Imaan Tamimi, Arian Jamasb, Marisa Dreher, Markus Pleijzier, Robert Turnbull, Nadiya Sharifi, Steven Calle, Andrew Dacks, Konrad Heinz, Kimberly Meechan, Aidan Smith, Najla Masoodpanah, Serene Dhawan, Peter Gibb, Corey Fisher, Claire Peterson, Jason Polsky, Tansy Yang, Katharina Eichler, Joseph Hsu, Irene Varela, Lucia Kmecova, Istvan Taisz, Jacob Ratliff, Kaylynn Coates, Anna Li, Marta Costa, Tyler Paterson, Claire Managan, Adam Heath, Katie Stevens, Jack Mccarty, Nora Forknall, Laurin Bueld, Neha Rampally, Zane Mitrevica, Kelli Fairbanks, Stanley Tran, Shada Alghailani, Quinn Vanderbeck, Lauren Warner, Henrique Ludwig, Jeremy Johnson and Levi Helmick, each of whom have contributed over 1,000 synapses. We principally thank the Wellcome Trust UK and US Drosophila Connectomics, Jefferis, Janelia Connectome Annotation Team, Bock, Preat, Wilson, Dacks, Hampel and Seeds groups for sharing their published and unpublished work in the Catmaid data set. Further, we thank Michael Reiser, Vivek Jayaraman, Arthur Zhao, Tatsuo Okubo, Jenny Lu and Kathi Eichler for identifying neuron matches in Catmaid which helped us build our ground truth data set and Mareike Selcho for helping to confirm our FlyWire identifications of octopaminegic neurons. Development and administration of the Catmaid tracing environment and analysis tools were funded in part by National Institutes of Health BRAIN Initiative grant 1RF1MH120679-01 to Davi Bock and Gregory Jefferis, with software development effort and administrative support provided by Tom Kazimiers (Kazmos GmbH) and Eric Perlman (Yikes LLC). We thank the flywire.ai community for allowing us to use their semi-automatic neuronal reconstructions, which took over 1,366,543 edits from human annotators to build from automatically reconstructed segments (Dorkenwald et al., 2022). FlyWire is supported by NIH BRAIN Initiative grant MH117815 to Murthy and Seung. Additional large-scale proofreading and infrastructure was supported by Wellcome awards 203261/Z/16/Z to Jefferis, and NIMH BRAIN Initiative award 1RF1MH120679-01 and NSF NeuroNex award DBI-2014862 to Bock and Jefferis. Those persons contributing more than 1,000 edits were: Doug Bland, Austin T Burke, Yijie Yin, Laia Serratosa Capdevila, Kyle Patrick Willie, Arti Yadav, Ryan Willie, Nash Hadjerol, Zairene Lenizo, Griffin Badalemente, J. Anthony Ocho, Shirleyjoy Serona, Dharini Sapkal, Anjali Pandey, Ben Silverman, Varun Sane, Zeba Vohra, regine salem, Mendell Lopez, J. Dolorosa, Imaan Tamimi, Chitra Nair, Dhwani Patel, Joshua Ban∼ez, Márcia Santos, Katharina Eichler, Shaina Mae Monungolh, Dustin Garner, Jay Gager, Joseph Hsu, Mark Larson, Bhargavi Parmar, Rey Adrian Candilada, Dhara Kakadiya, Alexandre Javier, Itisha Joshi, Michelle Pantujan, Irene Salgarella, James Hebditch, Philipp Schlegel, Kaushik Parmar, Darrel Jay Akiatan, Kendrick Joules Vinson, Marina Gkantia, Ariel Dagohoy, remer tancontian, Chan Hyuk Kang, Hane Two, Markus Pleijzier, Emil Kind, Olivia Sato, Yashvi Patel, Miguel Albero, Eva Munnelly, Katie Molloy, Christopher Dunne, Quinn Vanderbeck, Rashmita Rana, Merlin Moore, Lucia Kmecova, Alexis E Santana Cruz, Nadia Seraf, Usb, Claire McKellar, Monika Patel, Mareike Selcho, Greg Jefferis, Steven Calle, Siqi Fang, Arzoo Diwan, Sarah Morejohn, Christa Baker, Brian Reicher, Sangeeta Sisodiya, Tansy Yang, Paul Brooks, Selden, Marlon Blanquart, Hyungjun Choi, Celia D, Sanna Koskela, Joanna Eckhardt, Krzysztof Kruk, Wolf Huetteroth, Alisa Poh, Stefanie Hampel, Wes Murfin, Li Guo, Zhihao Zheng, Szi-chieh Yu, Jones, Farzaan Salman, Amalia Braun, Mark Lloyd Pielago, Nidhi Patel, Ben Gorko, Akanksha Jadia, Fernando J Figueroa Santiago and Urja Verma. Manuscripts describing the proofread and annotated FlyWire whole fly brain connectome are in preparation. We thank Marissa Sorek for assistance with community management and Ran Lu, Thomas Macrina, Kisuk Lee, J. Alexander Bae, Shang Mu, Barak Nehoran, Eric Mitchell, Sergiy Popovych, Jongpeng Wu, Zhen Jia, Manuel Castro, Nico Kemnitz, Dodam Ih for alignment and segmentation of the FAFB EM volume and registration to the original FAFB EM dataset. We thank Davi Bock, Eric Perlman and Stuart Berg and the flywire.ai project for helping us make our preliminary results available to the community. We thank Forrest Collman, Casey Schneider-Mizell, Chris Jordan, Derrick Brittain, Akilesh Haligeri for CAVE development and maintenance, and Kai Kuehner, Oluwaseun Ogedengbe, Jay Gager, Will Silversmith, Ryan Morey for Neuroglancer development, tools, and Codex development. Automatically detected mitochondria counts in the HemiBrain data set were pulled from neuPrint (Plaza et al., 2022) (https://connectome-neuprint.github.io/neuprint-python/docs/mitocriteria.html). We thank Laia Serratosa Capdevila for proofreading this manuscript.

## 6 Author Contributions

Given alphabetically by forename. Conceptualization, A.S.B., J.F. and P.S.; Methodology A.C., A.S.B., G.S.X.E.J., J.F., M.D. and N.E.; Software, A.C., A.S.B, G.S.X.E.J., J.F., M.D. and N.E. Validation, A.C., A.S.B, J.F., M.D., N.E., and P.S.; Formal Analysis, A.C., A.S.B, J.F., N.E. and Y.Y.; Investigation, A.S.B, J.F., M.D., N.E., Y.Y.; Resources, A.C., G.S.X.E.J., J.F., M.D., N.E. and P.S.; Data Curation, A.C., A.L., A.S.B, J.F., N.E., P.S., R.P., S.F.M., T.P., T.R., Y.Y. and V.H., Writing – Original Draft, A.S.B, G.S.X.E.J., J.F., N.E., and V.H.; Writing – Review & Editing, A.S.B, G.S.X.E.J., J.F., N.E. and V.H.; Visualization, A.C., A.S.B, J.F., N.E., and Y.Y.; Supervision, G.S.X.E.J. and J.F.; Project Administration, A.S.B., G.S.X.E.J. and J.F.; Funding Acquisition, A.S.B., G.S.X.E.J. and J.F.; We also credit as authors the key contributors to the FlyWire project. FlyWire Infrastructure and Management: S.D.; FlyWire Codex: A.M.; FlyWire Proofreader Training and Community Management: A.S., C.M., S-C.Y., K.E., M.C., P.S.; FlyWire Annotations and Cell Type Matching: P.S., K.E., A.S.B., G.J. FlyWire Leads: M.M. and S.S. Cambridge Lead: G.J.

## 7 Supplements

### 7.1 Supplemental data

#### 7.1.1 Supplemental data 1

A .csv file which gives each cell type (cell_type) we used to generate our ground truth data, the transmitter it is has been reported to express from the literature (nt), the study that made this report (citation) and the type of evidence it contained (evidence).

#### 7.1.2 Supplemental data 2

A .csv file which indicates each neuronal reconstruction used to generate our ground-truth data. Presynapses from each reconstruction were used. Columns provide a unique identifier (id) (which is the root ID for a flywire neuron, a skeleton ID for a CATMAID neuron and a bodyid for a hemibrain neuron), the designated transmitter from the literature (nt), and the data set from which the neuron comes (dataset). Full set of presynaptic locations available upon request. CATMAID reconstructions can be found on virtual fly brain: https://fafb.catmaid.virtualflybrain.org/. Flywire reconstructions can be found at: https://ngl.flywire.ai/. Hemibrain reconstructions can be found at: https://neuprint.janelia.org/?dataset=hemibrain.

#### 7.1.3 Supplemental data 3

A .csv file where each row is a single identified hemibrain reconstruction. Includes all such neurons used in our analysis. Columns provide the each neuron’s unique identifier (bodyid), morphological cell type (cell_type), neuron-level transmitter prediction (conf_nt), confidence score for that prediction (conf_ nt_p) and whether or not presynapses from this this neuron were included in our ground-truth data (in_ground_truth). In addition the percentage of synapse-level transmitter predictions for each transmitter class for each neuron is given (columns: acetylcholine, gaba, glutamate,dopamine, serotonin, octopamine).

#### 7.1.4 Supplemental data 4

A .csv file where each row is a single identified FlyWire reconstruction. Includes all such neurons used in our analysis. Columns provide the each neuron’s unique identifier (root id), morphological cell type where this is currently known (cell type), neuron-level transmitter prediction (conf_nt), confidence score for that prediction (conf_nt_p) and whether or not presynapses from this this neuron were included in our ground-truth data (in ground truth). In addition the percentage of synapse-level transmitter predictions for each transmitter class for each neuron is given (columns: acetylcholine, gaba, glutamate,dopamine, serotonin, octopamine). In addition, it contains some labels not available in the HemiBrain data set. This includes information on the hemisphere the neuron is in (side), its hemilineage designation in two nomenclatures (ito_lee_hemilineage, hartenstein_hemilineage). Because the root id for neurons is changing as neurons are edited in an active connectome project (Dorkenwald et al., 2022) we also supply the position of a point in the neuron to help identify it in FlyWire voxel space (pos_x, pos_y, pos_z) and an ID for the nucleus segmentation (nucleus_id).

#### 7.1.5 Supplemental data 5

A .zip archive containing .png files depicting each of the 161 brain hemilineages we have used from the FlyWire data set. Neurons in each hemilineage are coloured by their neuron-level transmitter predictions, hemilineage names given in the file name. Hemilineage labels for the FlyWire data set will be reported in an upcoming manuscript, Schlegel et al. 2023 (in prep).

